# HR-FRGS: A Novel Biomarker Discovery Protocol using Dual Layer Hypergraph Learning for NGS RNA Sequence Data

**DOI:** 10.64898/2026.09.21.753079

**Authors:** Manan Kumar Gupta, Mainak Paul, Soumen Kumar Pati

## Abstract

Biomarker discovery from high-dimensional RNA sequencing data remains challenging. Conventional methods such as differential expression analysis, pairwise protein-protein interaction networks, and weighted gene co-expression network analysis suffer from false-positive interactions, transitivity-driven noise, and arbitrary thresholds for feature selection. Addressing these limitations, this study proposed a novel framework, Hypergraph Regularised Fuzzy Rough Gene Selection (HR-FRGS), for knowledge-driven biomarker discovery. This pipeline integrates information-rich protein clustering with entropy filtering and PPI-pruned co-expression interactions to form a dual-layer hypergraph. A hypergraph-based scoring method that combines Random Walk with Restart diffusion and Hypergraph Betweenness Centrality is used to identify genes that are central in the biological network. Furthermore an autonomous fuzzy rough set selection eliminates arbitrary threshold dependency for gene selection. HR-FRGS was applied to four TCGA cancer cohorts: Lung Adenocarcinoma, Head and Neck Cancer, Kidney Clear Cell Carcinoma, and Colon Cancer, which reduced over 60,000 transcripts to compact biomarker panels. Validation using classical Machine Learning algorithms with cross-validation demonstrated that biomarker panels matched or exceeded the classification performance of the full transcriptome (AUC > 0.99, MCC > 0.94) and confirmed generalizability (MCC ≥ 0.893). Benchmarking against baseline methods and an ablation study showed the contribution of major parts of the algorithm. Functional enrichment analysis captures established pan-cancer hallmarks and cohort-specific oncogenic mechanisms, confirming the biological relevance of the identified genes.

## 1. Introduction

In recent years, cancer has remained a global health challenge, as over 20 million cases of cancer are diagnosed annually worldwide and will continue to rise in the number of cancer cases in the upcoming decades [1]. A biomarker is a measurable indicator used to distinguish between a cancerous state and a normal state in patients. Advances in next-generation sequencing technologies have revolutionised the field of oncology, enabling comprehensive profiling of tumour transcriptomes and the extraction of RNA-based biomarkers [2, 3]. The Cancer Genome Atlas (TCGA) repository is of significant importance in large-scale biomarker discovery studies, as it provides RNA-seq datasets across numerous cancer types, combining both tumour and healthy samples [4]. The expression of tumour-specific tissues or genes, as well as mutated and amplified genes, is useful for RNA-based biomarker discovery [2]. However, RNA-seq datasets like those in TCGA capture the expression of over 20,000 protein-coding genes, most of which are passenger genes, and very few are the main driver genes of cancer. The extraction of meaningful biological information from these high-dimensional data often requires specialised computational methods and a framework capable of distinguishing causal regulatory signals from correlational noise [5].

While differential gene expression analysis is one of the most widely used techniques in RNA-seq data, used to identify highly upregulated and downregulated genes between two or more samples, it alone is unable to identify driver genes or master regulators. Many genes show highly altered expression due to downstream effects. Most of them do not exhibit a regulatory role because many regulatory genes operate within a haemostatic range. Thus, selecting a gene solely based on log2fc is insufficient and can yield false results for important genes. To identify the driver genes from the differentially expressed genes (DEGs), the protein-protein interaction network is integrated [5]. To filter the degs based on their interactions and associate them with biological pathways, functional modules, and disease phenotypes, the PPIN (Protein-protein interaction network) and WGCNA (weighted gene co-expression network analysis) methods are often used in network biology. PPIN analysis facilitates the identification of hub genes based on degree and centrality metrics, as well as protein complexes or functional modules using topological clustering techniques. WGCNA, meanwhile, identifies co-expressed genes and clusters the genes based on similar gene expression patterns [6, 7, 8].

The PPIN provides static physical interaction between proteins. But the interaction data are generated from the high-throughput yeast two-hybrid (Y2H) experimental technique, which introduces many false-positive results and noise [9]. On the other hand, in WGCNA, co-expression suffers from transitivity: if Gene A regulates B, and B regulates C, A and C will act coherently, leading to a statistical edge between A and C even if they never physically interact [48]. The PPIN often exhibits a hairball effect (dense networks in which all nodes are almost connected, with no structure). Also, the PPIN exhibits pairwise interactions, or edges (dyads), in simple graph theory. But many proteins form complexes (polyads) and function as a single subunit consisting of three or more proteins [10]. Normal graph theory models often fail to capture generating results that are not always biologically true. In a clique, the network topology is distorted, artificially inflating centrality measures [11, 12].

Selecting a minimal subset of genes that serve as biomarkers is crucial, but the process can be biased in high-dimensional datasets. In RNA-seq data, many feature genes can be used to characterise the cancer, but they are redundant, do not improve diagnostic performance, and increase the risk of overfitting [13, 14, 15]. Selecting a subset of genes using an arbitrary threshold, such as a top-ranked selection strategy (e.g., selecting the top 50 or 100 genes), is a common choice in many research studies to reduce dimensionality. However, using an arbitrary threshold does not always yield a deterministic result. Different thresholds can lead to drastically different results and downstream effects. This leads to ignorance of the natural variability of the data, introducing researcher-specific bias. Also, relying on subjective thresholds often fails to yield robust, non-redundant results for clinically important biomarkers [16].

To address the above issues, this paper introduces a novel methodology that employs a hypergraph-integrated model with entropy pruning and fuzzy rough set gene selection. The proposed methodology follows a hierarchical filtering solution. We use graph entropy, an information-theoretic and structural complexity metric, to filter the protein cluster from PPIN. This approach eliminates high-entropy protein clusters and retains low-entropy, structurally stable protein complexes and dense functional modules [17]. The filtered PPI is then used for pruning the WGCNA network. The co-expressed edges, which are indirect, connect proteins at long distances and are not supported by protein-protein interactions; these edges are removed, resulting in a biologically supported correlation network. The hypergraph is used to overcome the limitations in the pairwise network model as it captures higher-order biological interactions, such as protein complex formation and feedback or feedforward loops [18]. The hypergraph is built as an integrated network where protein complexes are represented as structural hyperedges and co-expression modules as functional hyperedges. We then use hypergraph diffusion algorithms to find out genes that are central to the flow in the complex network [19, 20]. Finally, the genes, weighted by the hypergraph diffusion score, are selected using an autonomous fuzzy rough set method to avoid arbitrary gene selection and the risk of overfitting [13].

## 2. Related Work

This section reviews literature across three areas central to the proposed HR-FRGS framework: transcriptomic biomarker discovery, network-based gene prioritisation, and feature selection in high-dimensional genomic data.

Utilising population-scale transcriptomic profiling data, RNA sequencing has transformed biomarker discovery across many cancer types. While DESeq2-based differential gene expression analysis remains a standard filtering strategy for early-stage DEGs, it often captures the downstream effects rather than genes driving the cancer [3, 5]. Recent deep learning approaches, such as transformer-based models applied to transcriptomic data, have sought to improve gene prioritisation beyond statistical thresholding, yet they still lack biological interpretability. An arbitrary top-k gene selection strategy is used in most pipelines, leading to reproducibility issues [16, 21].

To move beyond univariate approaches, network-based methods such as protein-protein interaction and co-expression networks have generally been used. PPI-based networks are utilised for hub identification using degree and centrality measures, and for detecting functional modules and protein complexes using clustering algorithms such as MCODE [7, 10]. However, high-throughput yeast two-hybrid assays generate many false-positive interactions [9]. The WGCNA method builds gene-gene co-expression networks with a topological overlapping matrix and groups them using hierarchical clustering and soft thresholding [6, 7, 8]. WGCNA co-expression is prone to transitivity bias: indirect regulatory chains among genes yield spurious statistical edges lacking direct physical support [22]. The hypergraph-based models are used to capture complex higher-order biological interactions, such as feedback loops, thereby overcoming the limitations of conventional pairwise graph representations [18, 23]. To capture global regulatory signals from these models, a Random Walk with Restart is used, combined with a restart probability to identify influential genes in biologically meaningful subnetworks [20, 24].

In RNA-seq biomarker detection, feature selection and dimensionality reduction are crucial for downstream analysis. To overcome the arbitrary selection problem, fuzzy rough set theory can be applied to select a non-redundant, robust gene set, as it provides a principled, threshold-free measure of class dependency [25, 26]. Hu et al. have demonstrated that fuzzy similarity relations based on Gaussian kernels represent reliable uncertainty measurements for real-valued data sets [25], while Jensen and Shen have illustrated that dependency measurements in fuzzy rough sets surpass those of the classical rough set reduction technique in feature selection from noisy data with solid theoretical justification for genomic studies [27]. The greedy forward search based on marginal dependency gain can effectively approximate minimal sufficient features without setting an initial cardinality number [28]. These works collectively motivate an integrated framework for biomarker discovery that combines biologically grounded network scoring with fuzzy rough set-based autonomous feature selection, as proposed in HR-FRGS.

## 3. Methodology

The methodology section explains the overall architecture of HR-FRGS. The main objective of this framework is to apply an efficient pipeline to identify robust and biologically relevant biomarkers from noisy high dimensional transcriptomics data. Unlike traditional feature selection methods that generally fail to capture higher-order relationships between proteins or genes, our approach integrates biological domain knowledge with advanced hypergraph-based fuzzy-rough mathematical modelling to ensure both statistical significance and biological relevance.

The framework combines transcriptomic expression data and protein-protein structural interaction constraints to identify robust cancer biomarkers. The multi-stage framework is built using these three main concepts: (a) Dual-layer hypergraph construction using entropy-based pruning, (b) diffusion-based scoring on dual hypergraph, (c) Autonomous fuzzy-rough set gene selection.

### 3.1. Data acquisition and pre-processing

The transcriptomics data used in this study were obtained from the Xena browser for the GDC TCGA star count datasets. The four different cancer cohorts are used for gene expression quantification data [4, 29]. In RNA-seq data, many genes may have missing values or appear as zeros. These values are generally represented as missing and can be due to biological absence or technical artefacts. If these values are not handled correctly, they will produce false results in differential expression analysis [30]. From the expression matrices, genes with missing values exceeding 50% are removed [31]. After that, we handled the missing values using the K-nearest neighbour (KNN) imputation method, with k set to 10 [32, 31]. The physical protein-protein interactions data are retrieved from the STRING database for Homo sapiens (9606.protein.links.v12.0.txt.gz). Protein-protein interactions having a high confidence score above 0.7 are kept, and the rest are discarded [33, 34].

### 3.2. Dual-layer hypergraph construction

This section discusses the core methodology step by step, starting with data acquisition and pre-processing. followed by a detailed discussion of dual-layer hypergraph construction, diffusion-based scoring, and the subsequent integration of fuzzy rough set-based biomarker panel selection.

#### 3.2.1. Differential Expression Analysis

The first major step of this pipeline involves extracting a set of high-confidence candidate genes (*V*_*DEG*_) from the raw star count RNA matrix (*K*) from TCGA using the R/Bioconductor package DESeq2. The read counts *K*_*ij*_ of gene *i* and sample *j* are discrete and over-dispersed, and are modelled into a negative binomial distribution with mean *μ*_*ij*_ and dispersion *α*_*i*_.

To account for the sequencing depth and technical bias between each sample, the median-of-ratios method is used with a size factor, *s*_*j*_. The dispersion *α*_*i*_ is crucial for the statistical inference of differential expression between tumour and healthy replicates. The variability between the groups is also represented by *α*_*i*_, and the variance of each gene is calculated using *α*_*i*_ and is defined in eq. (1)

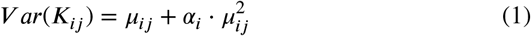

The resulting p-values are adjusted for multiple hypothesis testing using the Benjamini-Hochberg (BH) procedure to control the False Discovery Rate (FDR) [5, 35]. The resulting *V*_*DEG*_ is selected based on criteria often used in differential expression analysis [5, 36]:

- **Statistical Significance:** Adjusted p-value < 0.05
- **Magnitude of Effect:** | log _2_ Fold Change | >1.0 (representing a doubling or halving of expression)
- **Abundance:** Base mean expression > 10 counts (to remove noise)

The subset of *V*_*DEG*_ is used for downstream network formation as node input for PPIN and WGCNA.

#### 3.2.2. Protein-protein Network Construction Using Entropy Filtering

We used the *V*_*DEG*_ as nodes and the PPI from STRING as edges to build an undirected graph, *G* = (*V*_*DEG*_, *E*_*PPI*_). Even after using high-confidence thresholds for PPI, extracting centrality information from large PPINs becomes challenging, as they form messy, hairball-like structures that distort topology. To overcome this, the MCODE clustering algorithm is used to find dense topological communities or complexes. For MCODE, the fluff parameter is used to detect overlapping clusters, as a protein can participate in multiple functional complexes. This ensures that the proteins that act as bridges, participating in multiple important functional complexes, are accurately captured in the subsequent hypergraph-based scoring phase. The resulted clusters are represented as *C* = {*C*_1_, *C*_2_, *C*_3_, …, *C*_*K*_} where, *C*_*K*_ ⊆ *G*.

We use the structural information entropy as a metric to evaluate the quality of the clusters *C*. For each cluster, we calculate the normalised Shannon entropy *Ĥ* ∈ [0, 1] based on the internal degree distribution of the clusters. By applying this logic, we can identify dense, organised, and clique-like clusters (usually functional protein complexes) that have lower entropy than random, noisy, and disordered clusters [37]. We defined a subgraph for each cluster: *G*_*C*_ = (*V*_*C*_, *E*_*C*_). The degree (*d*_*i*_) for each node *i* is calculated in the clusters. The probability distribution (*P*_*k*_) is calculated using these degree values defined in eq. (2).

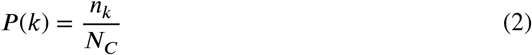

Where *n*_*k*_ is the number of nodes having degree *k*, and *N*_*C*_ is the total number of nodes in a cluster. We used the *P* (*k*) to calculate the Shannon entropy (*H*(*C*)) to measure the topological uncertainty in eq. (3).

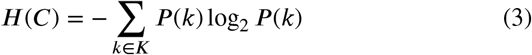

The resulting entropy *H*(*C*) is normalised in eq. (4) using the min-max method, where *H*(*C*) is divided by the highest possible entropy for the cluster *C*.

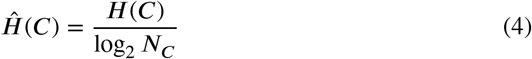

Where the entropy score *Ĥ* (*C*) ranges from 0 to 1, where low scores indicate a highly regular structure, such as a clique or a symmetric complex, in which nodes share uniform connectivity patterns. This represents a high-confidence functional module. We define an information-centric score in eq. (5) that accounts for high density, low entropy, and high average weight of the clusters.

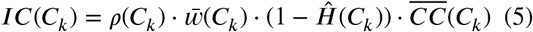

Where *ρ*(*C*_*k*_) is the density of each cluster, 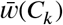 is the mean confidence score of the edges within the clusters, and 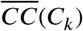 is the mean clustering coefficient. This *IC* serves as a primary weight for hypergraph construction. Finally, we filtered the cluster using the thresholds *ρ*(*C*_*k*_) ≥ 0.5 and *Ĥ*(*C*_*k*_) ≤ 0.8 to capture highly dense and ordered structures.

The entropy filtering eq. (2-5) were used to construct PPI network as shown in Algorithm 1.

#### 3.2.3. Weighted Gene Co-expression Network Analysis

Alongside the construction of the PPIN, we performed functional analysis using Weighted Gene Co-expression Network Analysis (WGCNA) on normalised expression data to identify Co-expressed gene modules. To explain the correlation strength between the genes, an adjacency matrix is built using eq. (6).

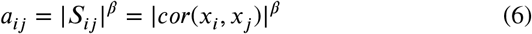

Where *a*_*ij*_ is the adjacency value between genes *x*_*i*_ and *x*_*j*_, *β* is the soft threshold parameter for scale-free topology (*R*_2_ = 0.85), and *S*_*ij*_ is the Pearson correlation of *x*_*i*_ and *x*_*j*_. We run WGCNA with a minimum module size of 20 [8, 38]. Subsequently, the adjacency matrix is converted into a topological overlap matrix (TOM). To reduce transitivity and noise, we implement a domain-aware pruning mechanism that integrates TOM similarity with the structural PPI scaffold. We define a structural penalty matrix (Ψ_*ij*_) in eq. (7) based on the geodesic distance *d*_*PPI*_ (*x*_*i*_, *x*_*j*_).

#### Algorithm 1

PPIEntropyFilter(*G, C, ρ*_min_,Ĥ_max_)

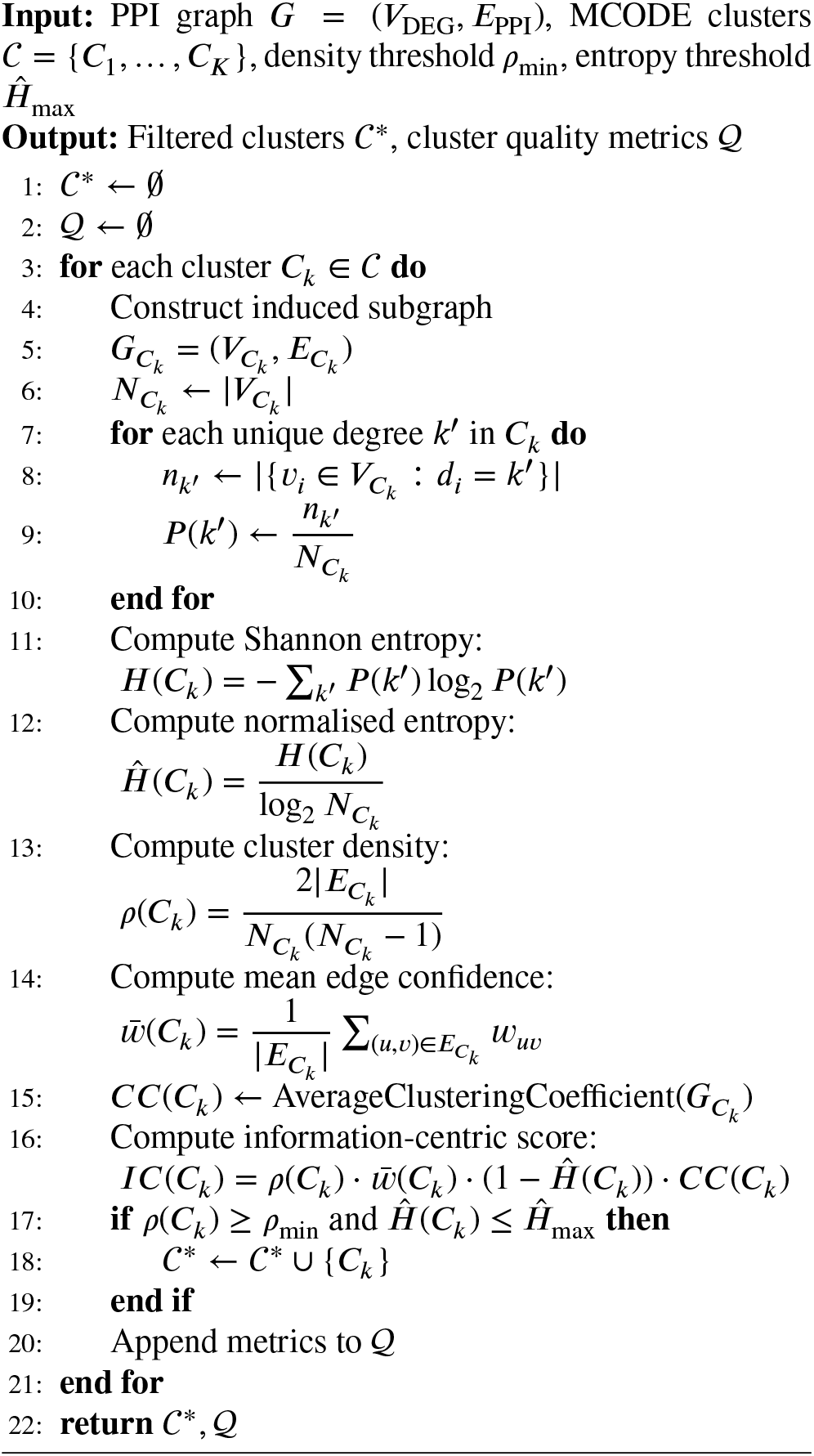

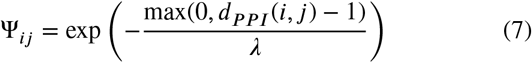

Where *λ* = 0.5. If the co-expressed genes lack a short physical interaction path, they are exponentially penalised. For example, if *d*_*PPI*_= 1 then Ψ_*ij*_ = 1, and if *d*_*PPI*_= 2 then Ψ_*ij*_ = 0.135. We performed hard pruning where *d*_*PPI*_≥ 3 and set Ψ_*ij*_ = 0 to remove co-expressed edges without biological support. The final pruned similarity matrix is defined using eq. (8).

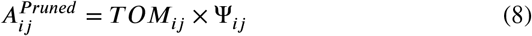

This approach efficiently filters the statistically co-expressed genes through a biologically grounded topological mask to highlight true causal drivers. To avoid over-pruning of highly co-expressed links, *A*_*ij*_ > 0.95 (High Confidence Threshold) is used, regardless of PPI distance. The construction of WGCNA TOM matrix and pruned adjacency matrix is shown in the Algorithm 2.

##### Algorithm 2

WGCNA Construction and Domain-Aware Pruning

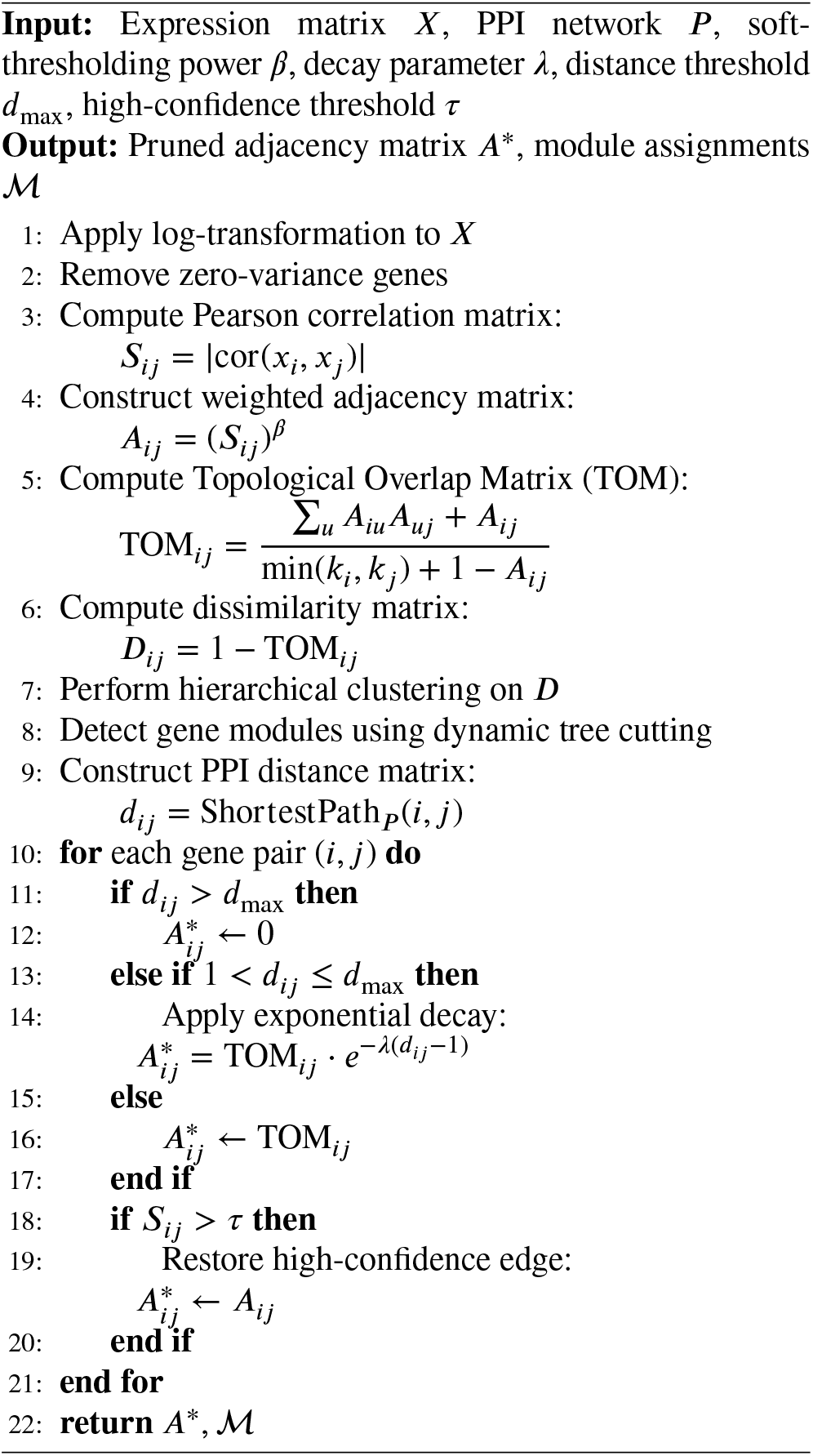

#### 3.2.4. Hypergraph Construction

Finally, using two filtered layers, filtered PPI (structural) and pruned adjacency (functional), we constructed an integrated hypergraph. For each structural hyperedge, each filtered PPI cluster (*C*_*k*_) is used and weighted using the *IC*(*C*_*k*_) score, whereas for the pruned adjacency, we applied distance-based hierarchical clustering to generate tight modules that capture genes with high correlation scores and weighted them using the module’s average adjacency. The total hypergraph is represented by an incidence matrix, *H* ∈ *R*^|*V* | ×| *E*|^ [18, 23]. The construction of dual-layer hypergraph is shown in Algorithm 3.

##### Algorithm 3

Dual-Weighted Hypergraph Construction

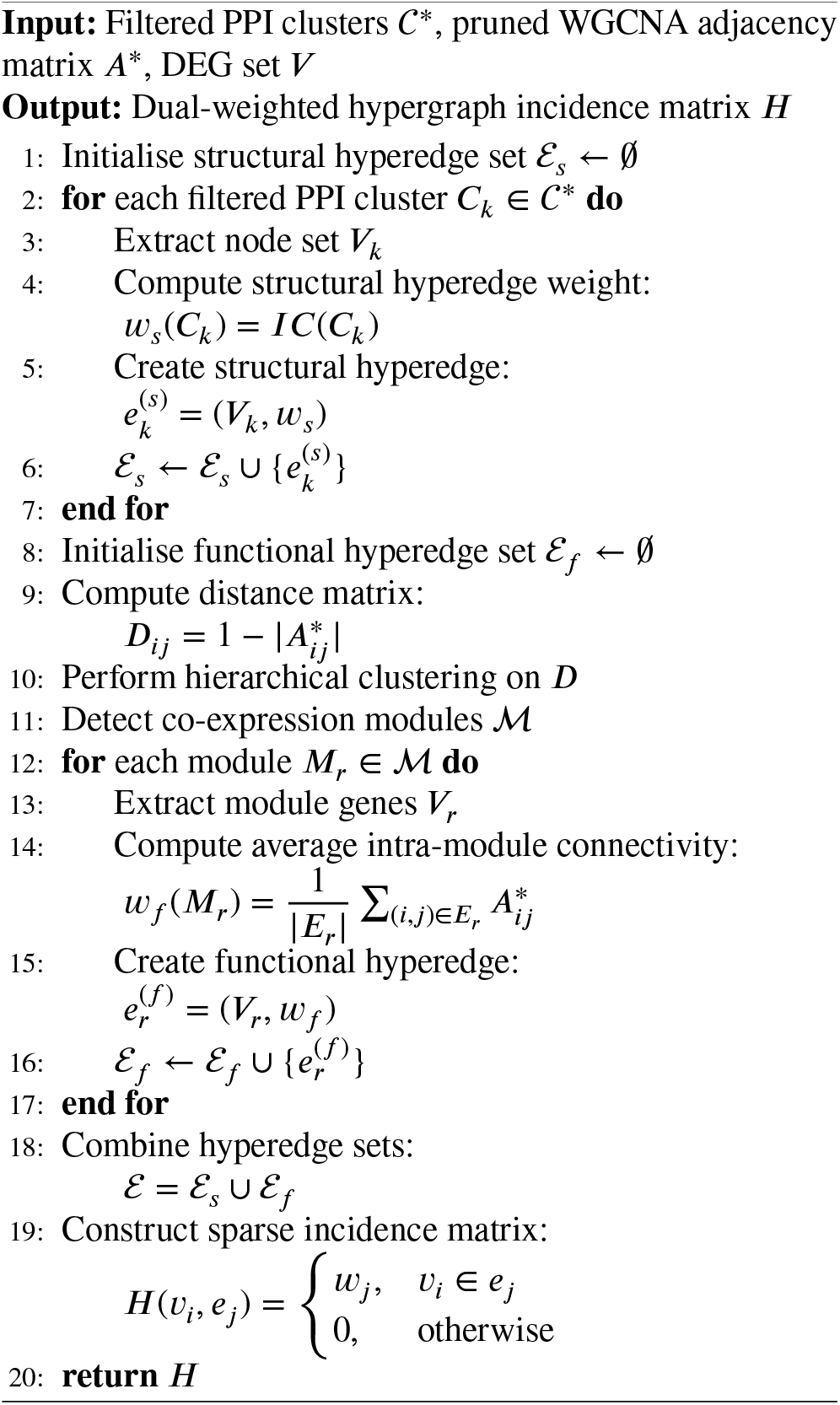

### 3.3. Diffusion-based Scoring on Dual-layer Hypergraph

For the scoring of genes using the dual-layer hypergraph first a transition matrix is built, then random walk with restart (RWR) and hypergraph betweeness centrality score is used to make a composite score for each gene.

#### 3.3.1. Transition Matrix Building

To identify the central genes in the biological flow, the simple degree centrality method is insufficient. We used hypergraph diffusion scoring by a random walk to observe the biological flow of information [24]. We calculate the two-step probability of the node *v*_*i*_ to *v*_*j*_. First, the probability of the node *v*_*i*_ to choose a hyperedge *e* is calculated, which is proportional to the hyperedge weight (*w*(*e*)). Then the probability of the walker travelling from the edge *e* to *v*_*j*_ is computed. To apply the random walk on the hypergraph, we construct a transition matrix *P*. The transition matrix *P* using eq. (9) captures the probability of information flow from the node *v*_*i*_ to node *v*_*j*_ across the hypergraph *H*.

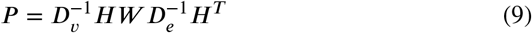

Where *W* is a diagonal matrix of hyperedge weights.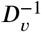 represents the inverse of the diagonal node degree matrix, which is the weighted sum of all possible incident hyperedges for each gene. *D*_*e*_ is the diagonal edge degree matrix, where each entry *D*_*e*_(*j, j*) represents the number of genes present in the edge *j*. The inverse of this matrix (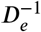) normalises the flow as it moves from an edge to a gene. This matrix-based approach is highly efficient, as it captures all possible communication paths and weights them according to their transition probabilities [19].

#### 3.3.2. Random Walk with Restart

The Random Walk with Restart (RWR) was applied in the transition matrix (*P*) to quantify the global influence of each gene within the dual-layer hypergraph [19, 20]. The RWR introduces a walker that iteratively explores the hyper-edges, modelling the biological flow of regulatory signals and calculating the total energy or influence with a restart probability [24]. This accumulates influence scores proportional to the connectivity and weight of the paths it traverses. The restart probability (*c*) ensures localising the walker in the influential region and prevents it from wandering in irrelevant regions. This is biologically sound as genes that are central to multiple high-confidence protein complexes and co-expression modules will naturally accumulate higher steady-state probability. The diffusion equation is defined in eq. (10).

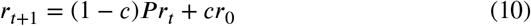

Where *r*_0_ is the initial seed vector, which has a uniform probability value of 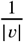. So, each gene begins with a uniform portion of total system energy. And *r*_*t*_ is the ranked vector at time *t. c* is the restart probability of the system (0.1). By iteratively calculating the transition matrix *P* with *r*_*t*_, the system converged at a particular threshold (10_*−*6_). When | |*r*_*t*+1_*− r*_*t*_| |_2_<10^*−*6^ is reached, the iteration stops, and the system reaches a steady state. The steady-state vector *r*_∞_represents the Diffusion Score [19]. Genes with high diffusion scores are not only locally connected but are globally influential across the entire biological interaction landscape, making them strong candidates for biomarkers [20].

#### 3.3.3. Hypergraph Betweenness Centrality

While the diffusion score captures reachability, to identify the major bottlenecks connecting different functional modules, the hypergraph betweeness centrality (HBC) is calculated by eq. (11). To overcome the complexity of calculating the centrality in a hypergraph, we performed a clique expansion [39]. This allows us to transform the incidence matrix *H* into a weighted undirected graph *G* = (*V*, *E*_*clique*_). The betweenness centrality *HBC*(*v*) is calculated by eq. (11).

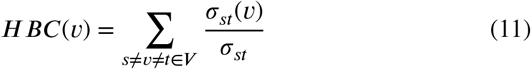

Where σ_*st*_ is the total number of weighted shortest hyperpaths between *s* and *t*, and σ_*st*_(*v*) is the number passing through *v*. The centrality vector is normalised to a range of [0, 1].

#### 3.3.4 Composite Biomarker Score

We define a Biomarker Potential Score (*BP S*) for each gene *v* by combining its diffusion score and centrality score as shown in eq. (12).

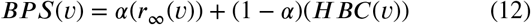

Where *α* is set to 0.5 to balance both scores. The higher *BP S* indicates a more central and influential gene.

### 3.4. Autonomous fuzzy-rough set gene selection

To select the genes that can classify samples into tumour and normal efficiently, without an arbitrary threshold that can bias the analysis, we used a fuzzy-rough set gene selection integrated with BPS value [13].

#### 3.4.1. Formulating Fuzzy-Similarity relationship

To integrate the *BP S* into the fuzzy-rough set model, we first normalised the *BP S* values into a vector *W* that has a sum of 1. First, we evaluate the weighted geometric distance between the sample *x*_*i*_ and sample *x*_*j*_ from the expression matrix for each gene *a* in the subset *B* using the eq. (13).

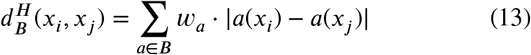

The distance is then used to construct a Gaussian kernel-based fuzzy similarity relationship using eq. (14) [25].

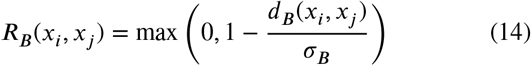

Where σ_*B*_ is the variance of the weighted distances. To keep the computational complexity minimal, all genes with *BP S* are clustered using the overlapping fuzzy c-means method. Then, for each cluster, the representative genes (those with the highest *IP D* score in the cluster) formed the gene candidate pool (*V*_*cand*_), which significantly reduces the search space for the final feature selection phase.

#### 3.4.2. Inner Product Dependency for Overlapping Classes

To evaluate the quality of a subset *B* of gene set *V*_*cand*_ and the fuzzy lower approximation 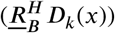 is calculated. It is then used to evaluate the classical dependency function, which calculates the size of the fuzzy positive region across all samples in the universe *U* using eq. (15).

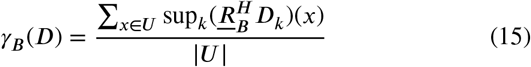

The γ_*B*_(*D*) only accounts for the maximum membership degree, but does not capture the class overlaps between tumour and normal samples. To resolve this problem, we use inner product dependency (*IP D*) using eq. (16).

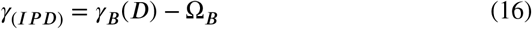

Where 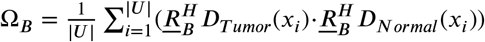 is the mean overlap of the decision classes. By using the *I P D*, we can penalise the gene set that has high duality for both classes [26]. To isolate the minimal subset *B* to maximise the *I P D* score, we used a greedy search algorithm. This algorithm iteratively adds a gene to the subset *B* and calculates the marginal dependency gain (Δγ). This acts as an exceptionally powerful multi-objective evaluation function.

#### 3.4.3. Autonomous Dynamic Halting via Discrete Curvature

However, the value of the Δγ decreases as the algorithm chooses the passenger genes. The value of the γ_(*IPD*)_ is plotted against the value of the |*S*| to form a curve that is concave downward. The optimal value of the gene set exists on the “Knee Point,” the key marker that indicates the point at which the systematic biological penalty for dimensionality (overfitting) begins to outweigh the statistical benefit of the greater class separability. The algorithm plots the values of the genes on the X-axis and the *IP D* values on the Y-axis to form a dependency curve. The values of each data point on the curve are normalised to determine the values for each data point. To mathematically determine the knee point on the curve, the algorithm calculates the discrete curvature κ on the normalised dependency curve. The data is discrete in nature. Therefore, the first derivative and the second derivative are calculated used in eq. (17) for each iteration *i*.

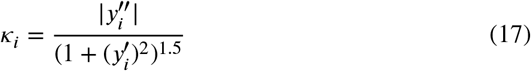

Where 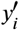 measures the marginal gain in *IP D* for each added gene and 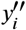measures the rate of diminishing returns (the “bend” in the^*i*^curve) and κ_i_ represent the curvature. The algorithm monitors the curvat^*i*^ure trend and autonomously triggers a halt at the global maximum of κ [28]. By halting exactly at the geometric knee point of the dependency curve, the HR-FRGS algorithm avoids traversing into the “tail” of the curve, which is mathematically composed almost entirely of overfitting, noise and redundant correlation structures. The whole structure of Fuzzy-rough gene selection is shown as Algorithm 4.

### 3.5. Internal validation

For evaluation of the performance of the potential biomarkers, we divided each cancer dataset into three categories based on the feature type: Total transcriptome (60660),

#### Algorithm 4

Hypergraph-Aware Fuzzy Feature Selection

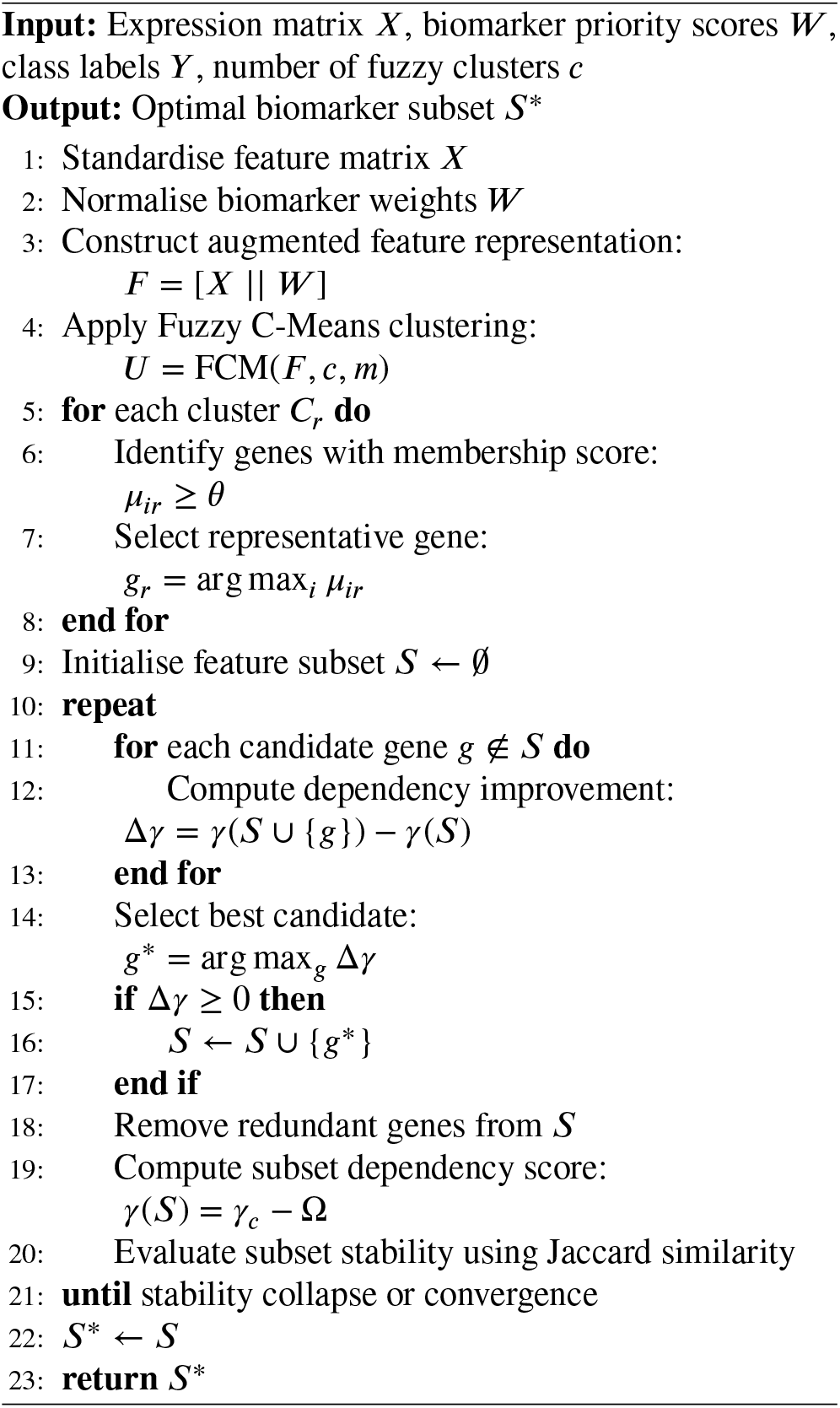

DEGs, and Potential biomarkers from FR gene selection and used them in a multi-metric performance comparison analysis. To perform a rigorous internal validation, three distinct supervised learning methods are used: Support Vector Machines (SVM), Random Forest (RF), and Extreme Gradient Boosting (XGBoost) with a repeated stratified k-fold cross-validation (5 folds, 10 repeats) [40, 41, 42]. In total, the 50 iterations help to measure the variance and stability of model performance, and stratification ensures the same ratio of different classes (“Tumour” and “Normal”) in each fold. The performance of classification is quantified with six evaluation metrics: Area Under Curve (AUC), Accuracy, F1-score, and Matthews Correlation Coefficient (MCC). The AUC was primarily used to measure the models’ ability to classify “Tumour” samples from “Normal” samples [43]. Accuracy and F1-score quantified models’ overall correctness and balance between the sensitivity and specificity, respectively [44]. MCC was used for more stringent evaluation as it accounts for all four quadrants of the confusion matrix and can produce reliable results for imbalanced TCGA datasets [45]. Together, these metrics are used to evaluate the classification power, clinical significance, and robustness of the selected biomarker panel and to compare with larger full transcriptome and DEG sets. Results are shown in Figures 1-5 and in Tables 2-3.

**Figure 1.**
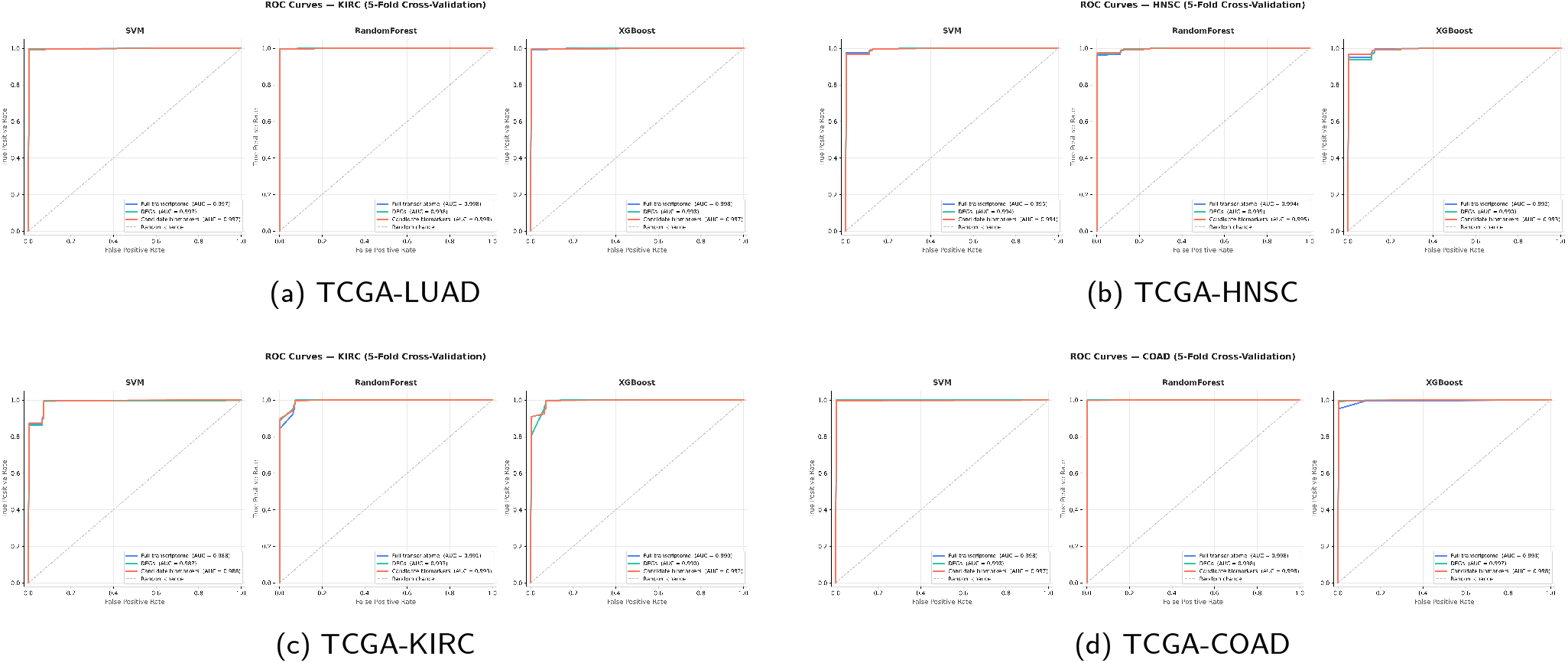
Receiver Operating Characteristic (ROC) curves for internal validation of the HR-FRGS biomarker panel across four TCGA cancer cohorts: (a) LUAD, (b) HNSC, (c) KIRC, and (d) COAD. Curves compare the discriminative performance of the Original transcriptome, DEG, and Biomarker feature sets using SVM, Random Forest, and XGBoost classifiers under repeated stratified 5-fold cross-validation.

**Figure 2.**
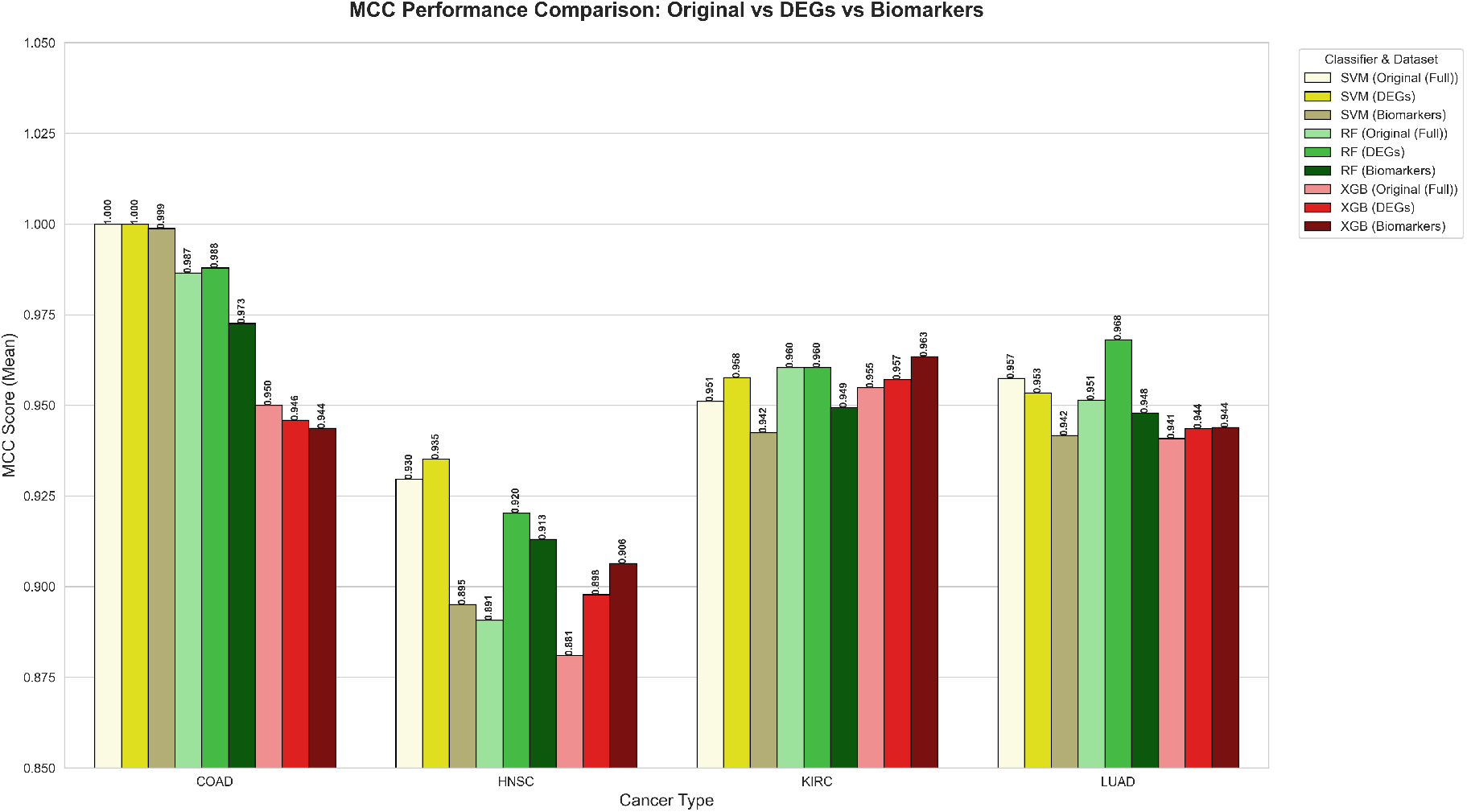
Comparison of Matthews Correlation Coefficient (MCC) across three classifiers (SVM, Random Forest, and XGBoost) for the Original, DEG, and HR-FRGS Biomarker feature sets on all four TCGA cancer cohorts (LUAD, HNSC, KIRC, COAD). Shaded bands represent variance across 50 cross-validation folds.

**Figure 3.**
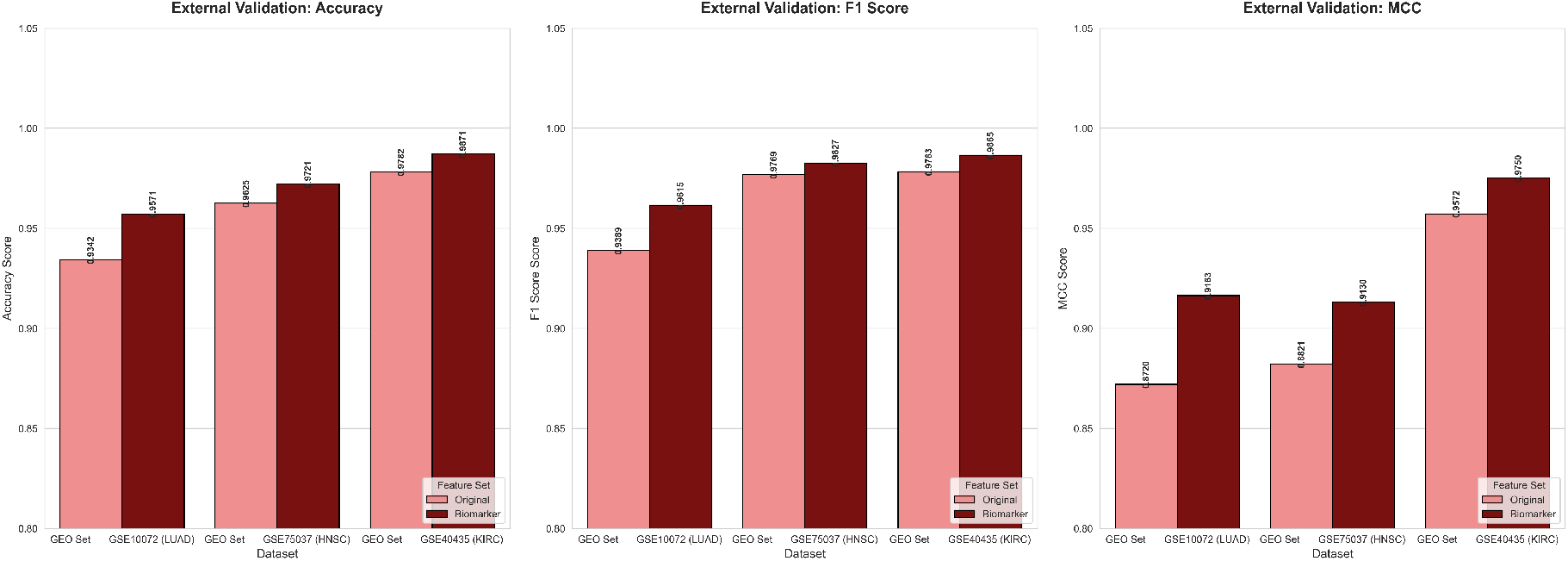
External validation classification performance of the HR-FRGS biomarker panel versus the full gene set using XGBoost across three independent GEO datasets: GSE10072 (LUAD), GSE75037 (HNSC), and GSE40435 (KIRC). Metrics shown include Accuracy, F1-score, AUC, and MCC.

**Figure 4.**
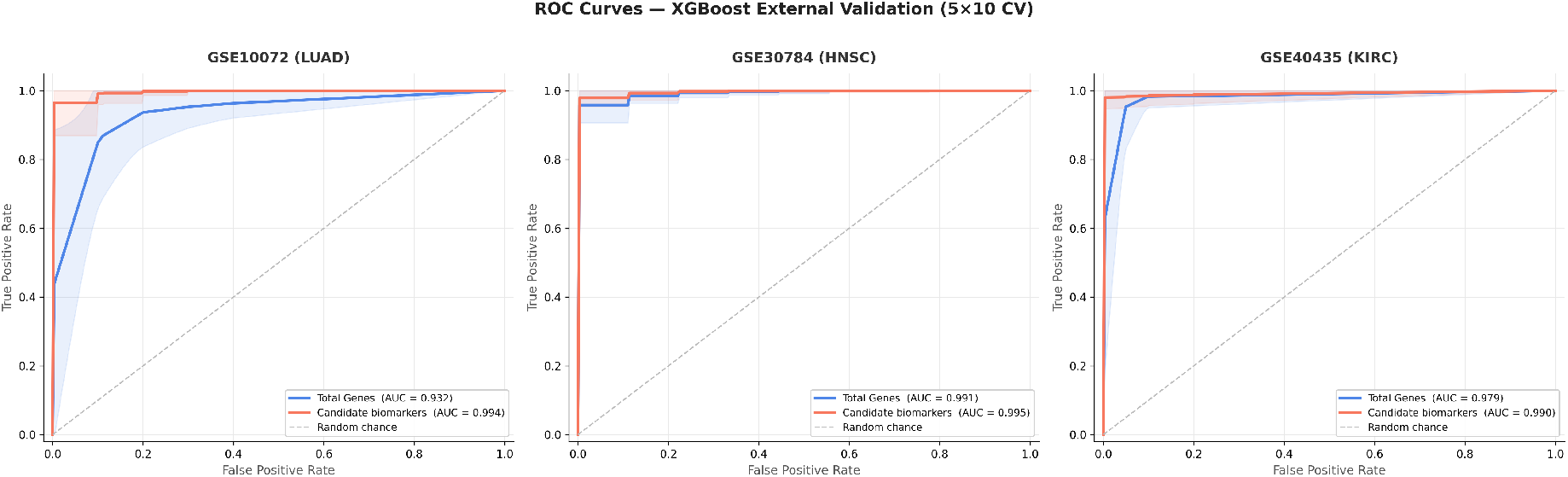
ROC-AUC curves comparing the HR-FRGS biomarker panel (red) and the full gene set (blue) on external GEO validation cohorts (GSE10072-LUAD, GSE75037-HNSC, GSE40435-KIRC) using the XGBoost classifier. Shaded regions represent 95% confidence intervals across 50 cross-validation folds.

**Figure 5.**
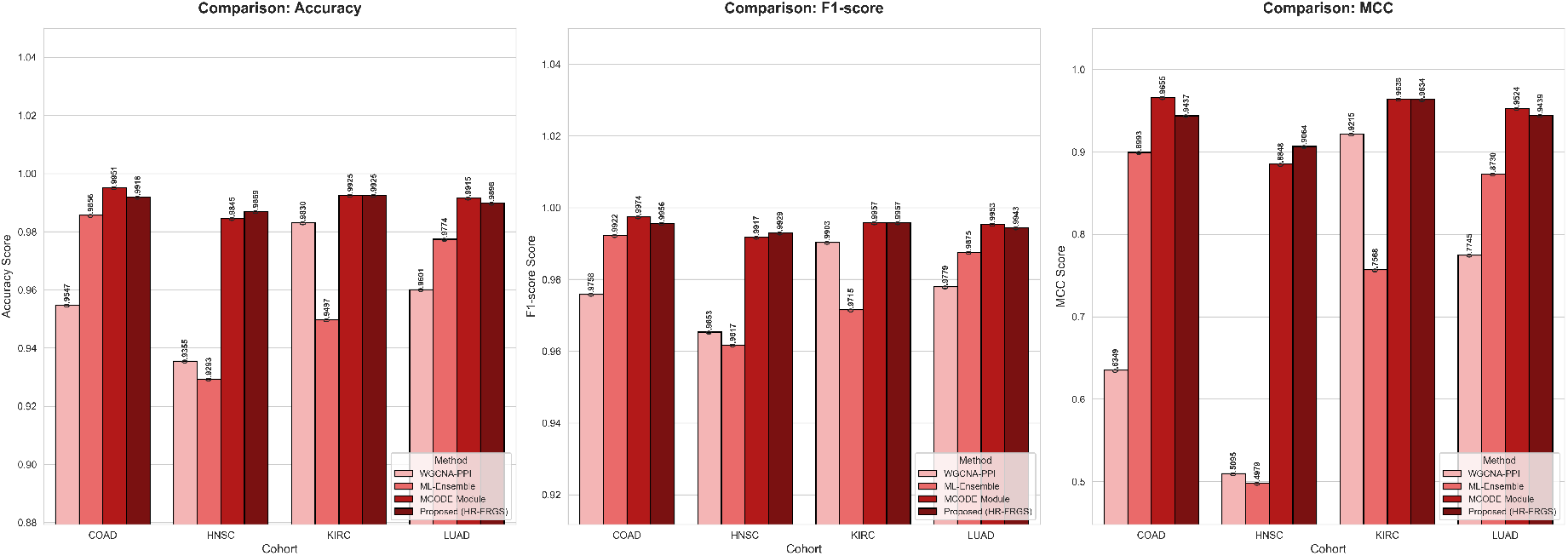
Bar chart comparison of classification performance (Accuracy, AUC, F1-score, and MCC) of HR-FRGS against three benchmark pipelines (WGCNA-PPI, ML-Ensemble, and MCODE-Module) using the XGBoost classifier across all four TCGA cancer cohorts (LUAD, HNSC, KIRC, COAD).

### 3.6. Validation of Potential Biomarker Panels in External Cohort

To evaluate the performance of the genes found by the HR-FRGS pipeline, validation on only the internal dataset is not sufficient. So, from the GEO (Gene Expression Omnibus) database, GSE10072, GSE75037, and GSE40435 datasets were extracted as external validation sets for LUAD, HNSC, and KIRC [46]. The LUAD cohort (GSE10072) comprised 58 tumour and 49 healthy samples; the HNSC cohort (GSE75037) included 184 tumour and 45 healthy samples; and the KIRC cohort (GSE40435) provided a perfectly balanced set of 101 tumour and 101 healthy samples. The external validation strategy was the same as the previous internal validation step, with each GEO dataset divided into two feature sets: a full gene set for baseline comparison and the target set containing identified common biomarkers. The classification strategy was kept the same as internal validation, where repeated stratified k-fold cross-validation (5 folds, 10 repeats) is used with XGBoost. To evaluate the performance, AUC, Accuracy, F1, and MCC metrics were used. MCC was treated as the primary evaluation metric given its robustness to class imbalance and its sensitivity to all four cells of the confusion matrix simultaneously [45]. Results are shown in Figures 6-7 and Table 4.

**Figure 6.**
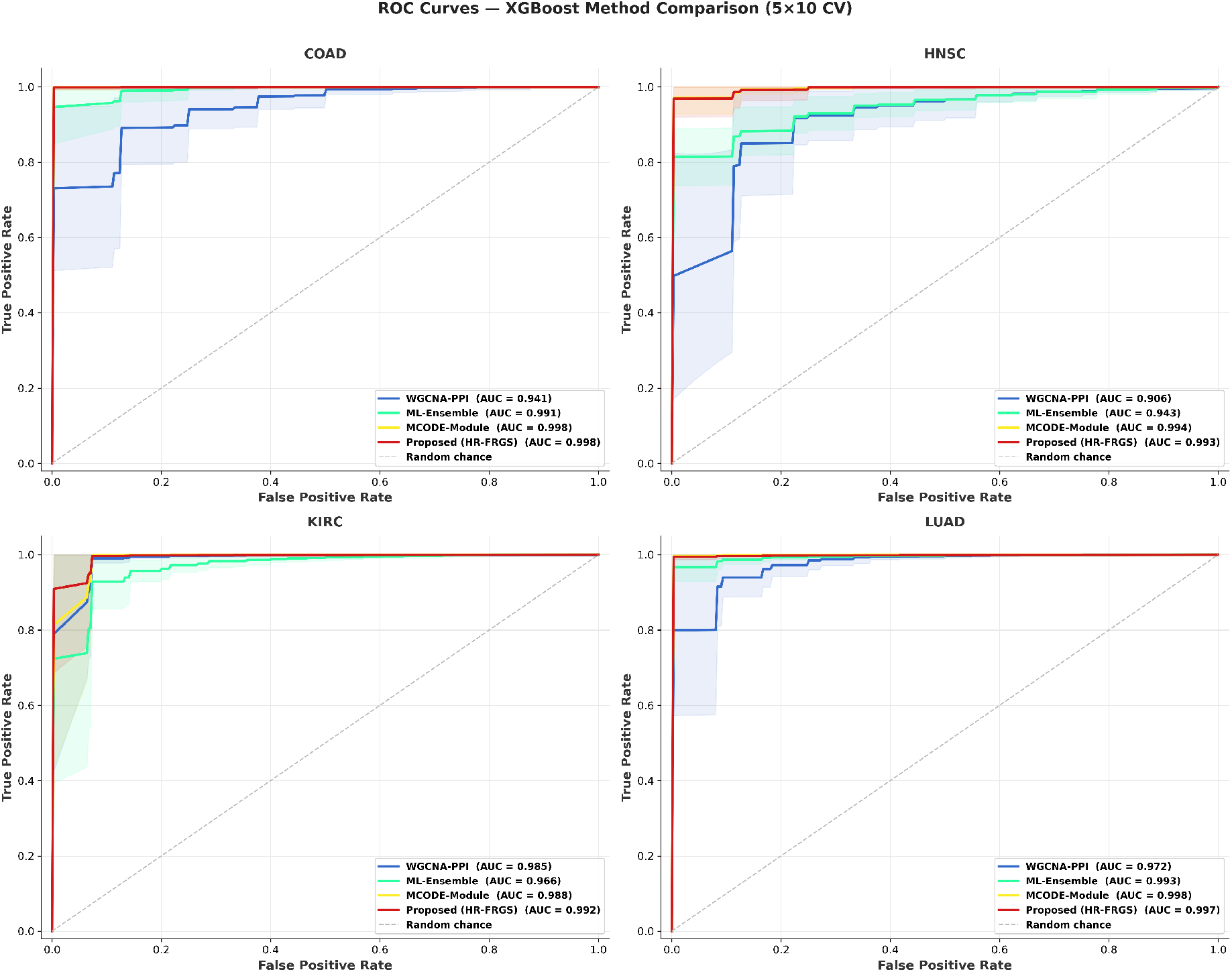
ROC-AUC curves comparing HR-FRGS and three benchmark biomarker discovery pipelines (WGCNA-PPI, ML-Ensemble, and MCODE-Module) using XGBoost across all four cancer cohorts (LUAD, HNSC, KIRC, COAD), illustrating the discriminative capacity of each method’s selected gene panel.

**Figure 7.**
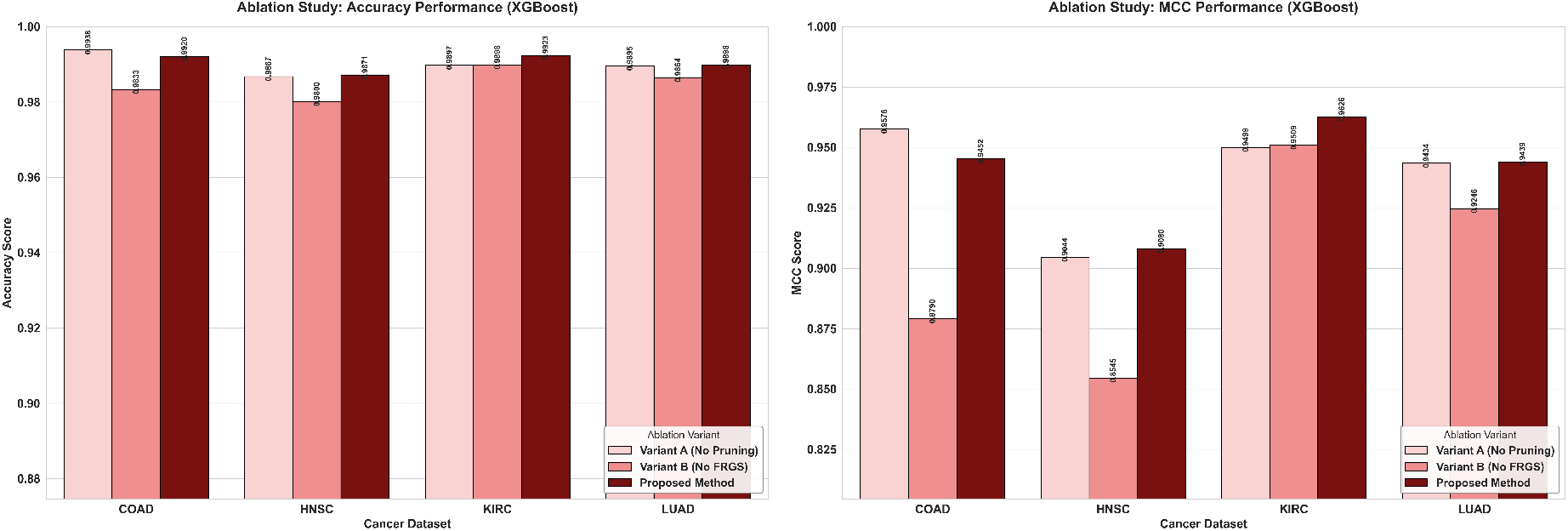
Ablation study results showing Accuracy and MCC performance of three HR-FRGS pipeline variants — Variant A (without entropy-based PPI cluster filtering), Variant (without fuzzy rough gene selection), and the complete HR-FRGS pipeline — using XGBoost on all four TCGA cancer cohorts.

### 3.7. Comparative Study and Performance Evaluation

The proposed pipeline HR-FRGS is benchmarked against three bioinformatics pipelines: WGCNA-PPI by Jia et al. [47], ML-Ensemble by Zhu et al. [48], and MCODE-Module by Kong et al. [49] with the same parameters, classification and metrics used for validation of external from the previous section. The three-benchmarking pipeline is also applied to all four of the cancer cohorts, and the number of biomarkers identified across the pipeline is shown in Table 6 and performance comparisons of each method are shown in Figure 8-9 and Table 5-6.

### 3.8. Ablation Study

The ablation study was conducted to evaluate the individual contributions of major sections in the pipeline to overall performance. The pipeline was divided into three different variants: Variant A with no entropy-based cluster filtering and Variant B with fuzzy-rough gene selection replaced by arbitrary gene selection. The performance of all results generated by these variants was evaluated against the original pipeline results using the same XGBoost classifier, metrics, and cross-validation strategy. The results are shown in Figure 10 and Table 7.

### 3.9. OncoKB validation and Functional Enrichment Analysis

The potential biomarkers are further validated using a curated cancer gene dataset from the OncoKB database [50]. The biomarkers were intersected with the OncoKB gene list to identify clinically relevant, established cancer-related genes as biological anchor seeds. To extract the remaining genes that are biologically relevant and can be evaluated as future cancer genes, a PPI neighbourhood expansion was performed using the STRING network with the same confidence threshold of ≥ 0.7. Genes that showed direct or one-hop physical interaction with any OncoKB seed gene in the PPIN were added to the computationally validated biomarker panel. This is a guilt-by-association approach that ensures that the added genes, which have a direct physical relationship with known oncogenic genes, are likely involved in the same functional pathway and mechanism in the relevant disease [51]. To characterise the functional and biological relevance of these genes, functional enrichment analyses were performed on the GO: BP, KEGG, and Reactome databases using the ClusterProfiler R package [52]. The over-representation analysis was performed for each cohort with an adjusted P-value < 0.05. The combined results of the enrichment analyses are shown as dot plots in Figure 8 for GO: BP, Figure 9 for KEGG, and Figure 10 for Reactome. The detailed term-level results in table format are supplied in the supplementary file.

## 4. Result

The above pipeline was applied in four distinct cancer datasets from Xena browser that are Lung Adenocarcinoma (LUAD) version 05-10-2024, Head and Neck Cancer (HNSC) version 05-10-2024, Kidney Clear Cell Carcinoma (KIRC) version 05-10-2024, and Colon Cancer (COAD) version 05-9-2024 [29]. All datasets initially have 60660 genes, but different numbers of samples: 589, 566, 610, and 514 for the cohorts LUAD, HNSC, KIRC, and COAD, respectively. After running the pipeline on these four cohorts, the number of potential biomarkers for each stage is shown in Table 1.

**Table 1.** Pipeline-level gene reduction summary across four TCGA cancer cohorts (LUAD, HNSC, KIRC, and COAD), showing the number of genes retained at each stage of the HR-FRGS framework from preprocessing through final biomarker selection.

| Datasets | Preprocessed genes | DEGs | Protein modules | WGCNA modules | Dual hyperedges | Final genes |
| --- | --- | --- | --- | --- | --- | --- |
| LUAD | 31857 | 6663 | 101 | 11 | 112 | 138 |
| HNSC | 29867 | 5136 | 75 | 5 | 80 | 282 |
| KIRC | 32969 | 7393 | 78 | 11 | 89 | 170 |
| COAD | 28670 | 5900 | 80 | 6 | 86 | 172 |

**Table 2.** Classification performance comparison (mean ± variance over 50-fold repeated stratified cross-validation) of three feature sets — full transcriptome (Original), differentially expressed genes (DEGs), and HR-FRGS biomarker panel — across four cancer cohorts (LUAD, HNSC, KIRC, COAD) using SVM, Random Forest, and XGBoost classifiers. Metrics reported: Accuracy, F1-score, and AUC.

| Cohort | Metric | SVM | Random Forest | XGBoost |
| --- | --- | --- | --- | --- |
| LUAD | Acc Original | 0.99185 ± 0.00005 | 0.99134 ± 0.00005 | 0.98947 ± 0.00009 |
| | Acc DEG | 0.99100 ± 0.00005 | 0.99423 ± 0.00003 | 0.98982 ± 0.00005 |
| | Acc Biomarker | 0.98880 ± 0.00007 | 0.99066 ± 0.00005 | 0.98981 ± 0.00007 |
| | F1 Original | 0.99545 ± 0.00002 | 0.99521 ± 0.00001 | 0.99418 ± 0.00003 |
| | F1 DEG | 0.99497 ± 0.00002 | 0.99680 ± 0.00001 | 0.99436 ± 0.00002 |
| | F1 Biomarker | 0.99374 ± 0.00002 | 0.99483 ± 0.00002 | 0.99435 ± 0.00002 |
| | AUC Original | 0.99905 ± 0.00000 | 0.99983 ± 0.00000 | 0.99922 ± 0.00000 |
| | AUC DEG | 0.99881 ± 0.00000 | 0.99974 ± 0.00000 | 0.99923 ± 0.00000 |
| | AUC Biomarker | 0.99833 ± 0.00000 | 0.99939 ± 0.00000 | 0.99860 ± 0.00001 |
| HNSC | Acc Original | 0.99011 ± 0.00009 | 0.98552 ± 0.00014 | 0.98392 ± 0.00011 |
| | Acc DEG | 0.99046 ± 0.00010 | 0.98922 ± 0.00008 | 0.98604 ± 0.00010 |
| | Acc Biomarker | 0.98445 ± 0.00012 | 0.98781 ± 0.00010 | 0.98692 ± 0.00008 |
| | F1 Original | 0.99466 ± 0.00003 | 0.99224 ± 0.00004 | 0.99136 ± 0.00003 |
| | F1 DEG | 0.99482 ± 0.00003 | 0.99421 ± 0.00002 | 0.99250 ± 0.00003 |
| | F1 Biomarker | 0.99156 ± 0.00004 | 0.99343 ± 0.00003 | 0.99295 ± 0.00002 |
| | AUC Original | 0.99620 ± 0.00003 | 0.99521 ± 0.00006 | 0.99325 ± 0.00017 |
| | AUC DEG | 0.99578 ± 0.00004 | 0.99639 ± 0.00004 | 0.99181 ± 0.00019 |
| | AUC Biomarker | 0.99519 ± 0.00002 | 0.99631 ± 0.00002 | 0.99513 ± 0.00008 |
| KIRC | Acc Original | 0.98951 ± 0.00005 | 0.99180 ± 0.00005 | 0.99066 ± 0.00007 |
| | Acc DEG | 0.99098 ± 0.00005 | 0.99180 ± 0.00006 | 0.99115 ± 0.00005 |
| | Acc Biomarker | 0.98770 ± 0.00006 | 0.98967 ± 0.00009 | 0.99246 ± 0.00007 |
| | F1 Original | 0.99405 ± 0.00002 | 0.99538 ± 0.00002 | 0.99473 ± 0.00002 |
| | F1 DEG | 0.99489 ± 0.00002 | 0.99538 ± 0.00002 | 0.99501 ± 0.00002 |
| | F1 Biomarker | 0.99302 ± 0.00002 | 0.99419 ± 0.00003 | 0.99575 ± 0.00002 |
| | AUC Original | 0.98953 ± 0.00032 | 0.99199 ± 0.00030 | 0.99162 ± 0.00027 |
| | AUC DEG | 0.98815 ± 0.00038 | 0.99453 ± 0.00016 | 0.99170 ± 0.00027 |
| | AUC Biomarker | 0.98974 ± 0.00031 | 0.99430 ± 0.00015 | 0.99354 ± 0.00019 |
| COAD | Acc Original | 1.00000 ± 0.00000 | 0.99805 ± 0.00002 | 0.99241 ± 0.00006 |
| | Acc DEG | 1.00000 ± 0.00000 | 0.99825 ± 0.00001 | 0.99162 ± 0.00009 |
| | Acc Biomarker | 0.99980 ± 0.00000 | 0.99611 ± 0.00006 | 0.99183 ± 0.00011 |
| | F1 Original | 1.00000 ± 0.00000 | 0.99895 ± 0.00000 | 0.99587 ± 0.00002 |
| | F1 DEG | 1.00000 ± 0.00000 | 0.99905 ± 0.00000 | 0.99544 ± 0.00003 |
| | F1 Biomarker | 0.99989 ± 0.00000 | 0.99790 ± 0.00002 | 0.99558 ± 0.00003 |
| | AUC Original | 1.00000 ± 0.00000 | 1.00000 ± 0.00000 | 0.99486 ± 0.00021 |
| | AUC DEG | 1.00000 ± 0.00000 | 1.00000 ± 0.00000 | 0.99905 ± 0.00001 |
| | AUC Biomarker | 1.00000 ± 0.00000 | 0.99990 ± 0.00000 | 0.99961 ± 0.00000 |

### 4.1. Internal validation

The performance of three datasets for each cohort across all classifier models is shown in Figures 1–2, and the detailed mean scores with total variance across 50 folds are given in Tables 2–5. The smaller candidate biomarker panels consistently matched or outperformed much larger transcriptome and DEG panels across all classifiers, cohorts, and metrics. In the LUAD cohort, the biomarker panel achieved very close to or superior scores, especially with XGBoost for the MCC metric (0.94392), indicating strong balanced discrimination. In the HNSC cohort, the observed performance of the biomarker panel is better than that of the other two datasets with the RF and XGBoost classifiers. Similarly, in the KIRC cohort, the biomarkers performed excellently with RF and XGBoost on the MCC metric (0.94941 and 0.96343, respectively), using only 170 genes. Lastly, the biomarkers of the COAD cohort achieved near-perfect classification scores with SVM and RF, and comparable performance with the XGBoost classifier. These results suggest that even after heavy dimensionality reduction (from 60,660 genes to 138–282 genes), the candidate biomarkers preserve the discriminative power of the full transcriptome and DEG sets, or even achieve higher power for classifying tumour and healthy samples across all four TCGA cohorts and three classifiers.

**Table 3.**
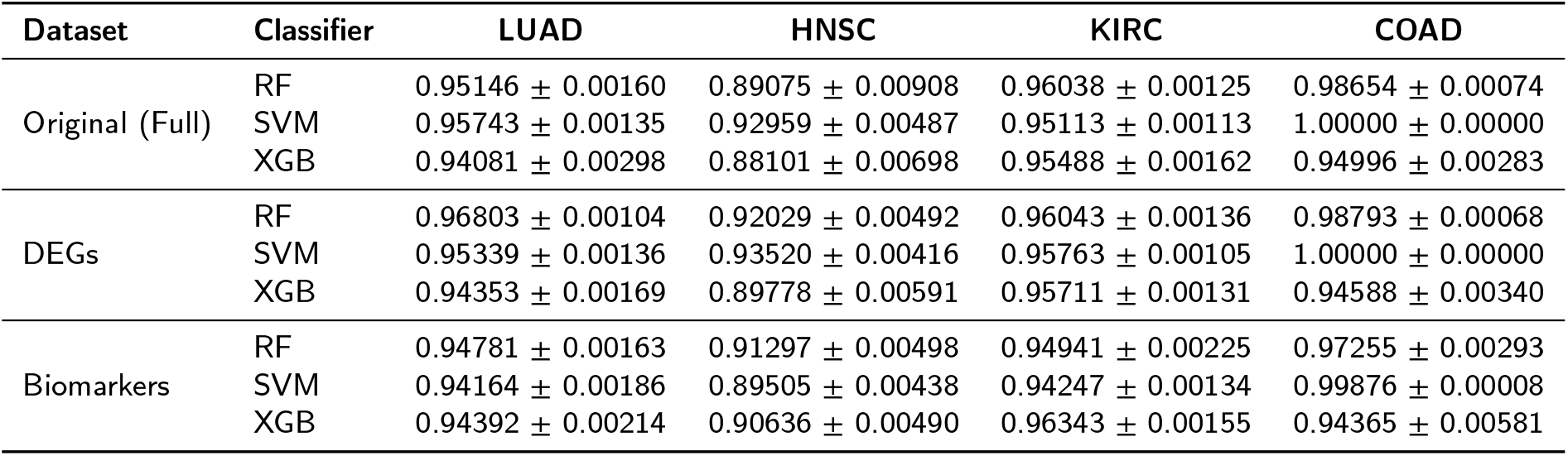
Matthews Correlation Coefficient (MCC) comparison (mean ± variance) across three classifiers (SVM, Random Forest, XGBoost) for the Original, DEG, and HR-FRGS Biomarker feature sets on all four TCGA cancer cohorts (LUAD, HNSC, KIRC, COAD).

| Dataset | Classifier | LUAD | HNSC | KIRC | COAD |
| --- | --- | --- | --- | --- | --- |
| Original (Full) | RF | 0.95146 ± 0.00160 | 0.89075 ± 0.00908 | 0.96038 ± 0.00125 | 0.98654 ± 0.00074 |
| | SVM | 0.95743 ± 0.00135 | 0.92959 ± 0.00487 | 0.95113 ± 0.00113 | 1.00000 ± 0.00000 |
| | XGB | 0.94081 ± 0.00298 | 0.88101 ± 0.00698 | 0.95488 ± 0.00162 | 0.94996 ± 0.00283 |
| DEGs | RF | 0.96803 ± 0.00104 | 0.92029 ± 0.00492 | 0.96043 ± 0.00136 | 0.98793 ± 0.00068 |
| | SVM | 0.95339 ± 0.00136 | 0.93520 ± 0.00416 | 0.95763 ± 0.00105 | 1.00000 ± 0.00000 |
| | XGB | 0.94353 ± 0.00169 | 0.89778 ± 0.00591 | 0.95711 ± 0.00131 | 0.94588 ± 0.00340 |
| Biomarkers | RF | 0.94781 ± 0.00163 | 0.91297 ± 0.00498 | 0.94941 ± 0.00225 | 0.97255 ± 0.00293 |
| | SVM | 0.94164 ± 0.00186 | 0.89505 ± 0.00438 | 0.94247 ± 0.00134 | 0.99876 ± 0.00008 |
| | XGB | 0.94392 ± 0.00214 | 0.90636 ± 0.00490 | 0.96343 ± 0.00155 | 0.94365 ± 0.00581 |

**Table 4.** External validation performance (mean ± variance, 50-fold repeated stratified 5-fold cross-validation) of the full gene set and HR-FRGS biomarker panel using XGBoost on three independent GEO cohorts: GSE10072 (LUAD), GSE75037 (HNSC), and GSE40435 (KIRC). Metrics reported: Accuracy, F1-score, AUC, and MCC.

| Metric | Set | GSE10072 (LUAD) | GSE75037 (HNSC) | GSE40435 (KIRC) |
| --- | --- | --- | --- | --- |
| Accuracy | Original | 0.9342 ± 0.0002 | 0.9625 ± 0.0001 | 0.9782 ± 0.0001 |
| | Biomarker | 0.9571 ± 0.0002 | 0.9721 ± 0.0000 | 0.9871 ± 0.0000 |
| F1 | Original | 0.9389 ± 0.0002 | 0.9769 ± 0.0000 | 0.9783 ± 0.0001 |
| | Biomarker | 0.9615 ± 0.0002 | 0.9827 ± 0.0000 | 0.9865 ± 0.0000 |
| AUC | Original | 0.9419 ± 0.0003 | 0.9930 ± 0.0000 | 0.9782 ± 0.0001 |
| | Biomarker | 0.9965 ± 0.0000 | 0.9955 ± 0.0000 | 0.9917 ± 0.0000 |
| MCC | Original | 0.8720 ± 0.0007 | 0.8821 ± 0.0009 | 0.9572 ± 0.0003 |
| | Biomarker | 0.9163 ± 0.0008 | 0.9130 ± 0.0006 | 0.9750 ± 0.0001 |

### 4.2. Validation of Potential Biomarker Panels in External Cohort

From Table 4 and Figurers 3-4, the validation of biomarkers in the external GSE10072(LUAD) cohort using XG-Boost shows that while the full gene set achieved Accuracy = 0.934, F1 = 0.939, AUC = 0.942, and MCC = 0.872, the biomarker panel outperformed it across all metrics by achieving Accuracy = 0.957, F1 = 0.962, AUC = 0.997, and MCC = 0.916. Using biomarkers, XGBoost achieved an MCC of 0.044, suggesting that the biomarker panel captures more discriminative information and reduces noise. In the GSE75037 (HNSC) cohort, although both gene sets showed stable performance, the biomarker panel achieved an accuracy of 0.972, an F1 score of 0.983, an AUC of 0.996, and an MCC of 0.913. The 0.033 gain in MCC using the biomarkers confirmed strong generalisation of the HNSC biomarker panel to independent data, despite the molecular heterogeneity of this cancer type. Also, in the case of the KIRC external dataset (GSE40435), both of the gene sets performed near-perfectly, but still the biomarkers consistently outperformed: Accuracy = 0.987, F1 = 0.987, AUC = 0.992, and MCC = 0.975, gaining 0.0089 higher Accuracy and 0.0178 higher MCC than the full gene set. The ROC curves in Figure 7 further explain these findings, with the biomarker panel (red) achieving higher AUC than the full gene set (blue) across all three cohorts and displaying consistently narrower confidence bands. For the GSE10072 dataset, the difference in AUC (0.994 vs 0.932) confirms that the biomarkers improved the classifier’s confidence in this cohort. These patterns confirm that the HR-FRGS-derived biomarker panels are not TCGA-specific artefacts but rather biologically important, robust molecular signatures that retain and even enhance their discriminative power on entirely independent datasets.

### 4.3. Comparative Benchmarking and Dimensionality Analysis

The comparative analysis from Table 5-6 and Figure 5-6 reveals distinct results of the four methods. The WGCNA-PPI and ML-Ensemble methods are very restrictive in identifying a number of biomarkers of 10 or fewer. The extreme reduction and arbitrary selection can introduce an information bottleneck and eliminate important regulatory genes necessary for understanding complex tumour microenvironments. On the other hand, the MCODE-Module approach identified 321-455 genes, likely introducing passenger genes that can contribute to noise and overfitting.

**Table 5.** Comparison of biomarker panel dimensionality across four cancer cohorts (LUAD, HNSC, KIRC, COAD) for HR-FRGS and three benchmark pipelines: MCODE-Module, ML-Ensemble, and WGCNA-PPI.

| Dataset | HR-FRGS | MCODE-Module [47] | ML-Ensemble [48] | WGCNA-PPI [49] |
| --- | --- | --- | --- | --- |
| LUAD | 138 | 391 | 8 | 10 |
| HNSC | 282 | 332 | 4 | 10 |
| COAD | 172 | 321 | 4 | 10 |
| KIRC | 170 | 455 | 5 | 10 |

**Table 6.** Comparative benchmarking of HR-FRGS against three state-of-the-art biomarker discovery pipelines (WGCNA-PPI, ML-Ensemble, and MCODE-Module) using XGBoost classification on all four TCGA cancer cohorts. Mean performance is reported across Accuracy, AUC, F1-score, and MCC metrics.

| Metric | Method | COAD | HNSC | KIRC | LUAD |
| --- | --- | --- | --- | --- | --- |
| Accuracy | WGCNA-PPI | 0.9547 ± 0.0002 | 0.9355 ± 0.0004 | 0.9830 ± 0.0001 | 0.9601 ± 0.0002 |
| | ML-Ensemble | 0.9856 ± 0.0001 | 0.9293 ± 0.0004 | 0.9497 ± 0.0004 | 0.9774 ± 0.0002 |
| | MCODE-Module | 0.9951 ± 0.0001 | 0.9845 ± 0.0001 | 0.9925 ± 0.0000 | 0.9915 ± 0.0001 |
| | Proposed (HR-FRGS) | 0.9918 ± 0.0001 | 0.9869 ± 0.0001 | 0.9925 ± 0.0001 | 0.9898 ± 0.0001 |
| AUC | WGCNA-PPI | 0.9421 ± 0.0016 | 0.9072 ± 0.0031 | 0.9862 ± 0.0004 | 0.9734 ± 0.0006 |
| | ML-Ensemble | 0.9927 ± 0.0002 | 0.9440 ± 0.0005 | 0.9673 ± 0.0008 | 0.9943 ± 0.0000 |
| | MCODE-Module | 0.9998 ± 0.0000 | 0.9956 ± 0.0000 | 0.9896 ± 0.0004 | 0.9996 ± 0.0000 |
| | Proposed (HR-FRGS) | 0.9996 ± 0.0000 | 0.9951 ± 0.0001 | 0.9935 ± 0.0002 | 0.9986 ± 0.0000 |
| F1-score | WGCNA-PPI | 0.9758 ± 0.0001 | 0.9653 ± 0.0001 | 0.9903 ± 0.0000 | 0.9779 ± 0.0001 |
| | ML-Ensemble | 0.9922 ± 0.0000 | 0.9617 ± 0.0001 | 0.9715 ± 0.0001 | 0.9875 ± 0.0000 |
| | MCODE-Module | 0.9974 ± 0.0000 | 0.9917 ± 0.0000 | 0.9957 ± 0.0000 | 0.9953 ± 0.0000 |
| | Proposed (HR-FRGS) | 0.9956 ± 0.0000 | 0.9929 ± 0.0000 | 0.9957 ± 0.0000 | 0.9943 ± 0.0000 |
| MCC | WGCNA-PPI | 0.6349 ± 0.0314 | 0.5095 ± 0.0292 | 0.9215 ± 0.0019 | 0.7745 ± 0.0069 |
| | ML-Ensemble | 0.8993 ± 0.0053 | 0.4979 ± 0.0224 | 0.7568 ± 0.0081 | 0.8730 ± 0.0055 |
| | MCODE-Module | 0.9655 ± 0.0032 | 0.8848 ± 0.0066 | 0.9638 ± 0.0008 | 0.9524 ± 0.0021 |
| | Proposed (HR-FRGS) | 0.9437 ± 0.0058 | 0.9064 ± 0.0049 | 0.9634 ± 0.0016 | 0.9439 ± 0.0021 |

**Table 7.** Ablation study results comparing XGBoost classification performance (Accuracy and MCC, mean ± variance) across three HR-FRGS pipeline variants — Variant A (without entropy-based cluster filtering), Variant B (without fuzzy rough gene selection), and the full HR-FRGS pipeline — on all four cancer cohorts (LUAD, HNSC, KIRC, COAD).

| Metric | Variant | LUAD | HNSC | KIRC | COAD |
| --- | --- | --- | --- | --- | --- |
| Accuracy | Variant A (No Pruning) | 0.9895 ± 0.0001 | 0.9867 ± 0.0001 | 0.9897 ± 0.0001 | 0.9938 ± 0.0001 |
| | Variant B (No FRGS) | 0.9864 ± 0.0001 | 0.9800 ± 0.0002 | 0.9898 ± 0.0001 | 0.9833 ± 0.0001 |
| | Proposed (HR-FRGS) | 0.9898 ± 0.0001 | 0.9871 ± 0.0001 | 0.9923 ± 0.0001 | 0.9920 ± 0.0001 |
| MCC | Variant A (No Pruning) | 0.9434 ± 0.0023 | 0.9044 ± 0.0058 | 0.9499 ± 0.0016 | 0.9576 ± 0.0035 |
| | Variant B (No FRGS) | 0.9246 ± 0.0026 | 0.8545 ± 0.0093 | 0.9509 ± 0.0017 | 0.8790 ± 0.0084 |
| | Proposed (HR-FRGS) | 0.9439 ± 0.0021 | 0.9080 ± 0.0045 | 0.9626 ± 0.0017 | 0.9452 ± 0.0057 |

The comparative benchmarking performances reveal that in both LUAD and KIRC cohorts, the proposed HR-FRGS pipeline achieved a very high Accuracy score > 0.98 and MCC score > 0.94 across all classifiers, comparable to the MCODE-Module approach despite a substantially smaller gene set (138 vs 391 genes in LUAD; 170 vs 455 in KIRC) and significantly higher than ML-Ensemble and WGCNA-PPI approach, suggesting more robust performance with an optimal number of genes. Both HR-FRGS and MCODE-Module methods significantly outperformed ML-Ensemble (MCC: 0.838-0.873) and WGCNA-PPI (MCC: 0.765-0.775) in LUAD. This suggests that very restricted gene selection can cause an information bottleneck despite a lower number of feature sets. In the HNSC dataset, the HR-FRGS method clearly outperformed the other methods across all classifiers and metrics, with MCC ranging from 0.895 to 0.913, highlighting the stability of this approach in a complex heterogeneous dataset. In the COAD, all methods except WGCNA-PPI performed well. HR-FRGS maintained MCC = 0.944 and Accuracy = 0.9918 with 172 genes, comparable discriminative power with MCODE-Module, but with a more compact feature set. The ROC-AUC curves in Figure 9 further corroborate these findings, with HR-FRGS achieving AUC = 0.999 (COAD), 0.995 (HNSC), 0.994 (KIRC), and 0.999 (LUAD), closely matching MCODE-Module while substantially exceeding ML-Ensemble and WGCNA-PPI across all cohorts. Collectively, these results establish that HR-FRGS achieves the optimal balance between feature parsimony and classification robustness across all four cancer types.

### 4.4. Ablation Study

The ablation results using the XGBoost classifier are presented in Table 7 and Figure 7. Variant B consistently underperformed across all cohorts and classifiers. Variant B dropped from 0.944 to 0.925 (−0.019) in LUAD and from 0.945 to 0.879 (−0.066) in COAD, respectively. Also, in HNSC, the MCC score dropped for XGBoost. This suggests that without IPD-based class-overlap penalisation, feature selection can fail to identify the minimal discriminative subset across heterogeneous cancer types. Variant A also showed consistently moderate performance, particularly in HSNC for MCC (0.908 to 0.904), confirming that, without entropy filtering, unpruned clusters can introduce functional noise.

### 4.5. OncoKB validation and Functional Enrichment Analysis

In the resulting gene panel of four cohorts: LUAD, HNSC, KIRC, and COAD, the numbers of upregulated genes are 26, 122, 32, 48 and downregulated genes are 11, 32, 6, 15, respectively. In all four cancer cohorts, cell cycle regulation and DNA-related processes are consistently enriched biological phenomena. The GO: BP analysis shows that DNA recombination and protein-DNA complex assembly are significantly enriched in LUAD, whereas chromosome segregation, chromosome organisation, and DNA metabolic process are enriched in both HNSC and COAD cohorts, with high gene ratios and low adjusted p-values (Figure 8). The GO: BP result complements both KEGG and Reactome results, as the cell cycle emerged as highly significant in KEGG for both LUAD and HNSC (Figure 9). Also, in Reactome analysis across the three cohorts, LAUD, HNSC, and COAD show cell cycle, cell cycle checkpoint, and DNA replication as dominant terms, with high gene ratios (0.3-0.5) and exceptionally low adjusted p-values ranging from from 10^*−*22^ to 10^*−*29^ in HNSC in HNSC (Figure 10). In the three independent cohorts, cell cycle machinery and related processes are consistently enriched, capturing the core oncogenic process and cancer development rather than cohort-specific noise. The role of cell cycle dysregulation has a well-established role for cancer as a pan-cancer hallmark [53]. Chromatin organisation and epigenetic regulation emerged as the second shared phenomena in LAUD, HNSC and COAD. GO: BP terms such as epigenetic regulation of gene expression and heterochromatin formation were enriched in COAD, the same as chromosome segregation, chromosome organisation for HNSC, and protein-DNA complex for LUAD. Reactome results correlate this with enrichment of HDACs, which deacetylate histones; HATs, which acetylate histones; Nucleosome assembly; Deposition of new CENPA-containing nucleosomes at the centromere in COAD; and NoRC negatively regulates rRNA expression and Negative epigenetic regulation of rRNA expression in LUAD. These results are biologically consistent with the known roles of histone modifications and chromatin remodelling in transcriptional reprogramming during tumorigenesis [54].

**Figure 8.**
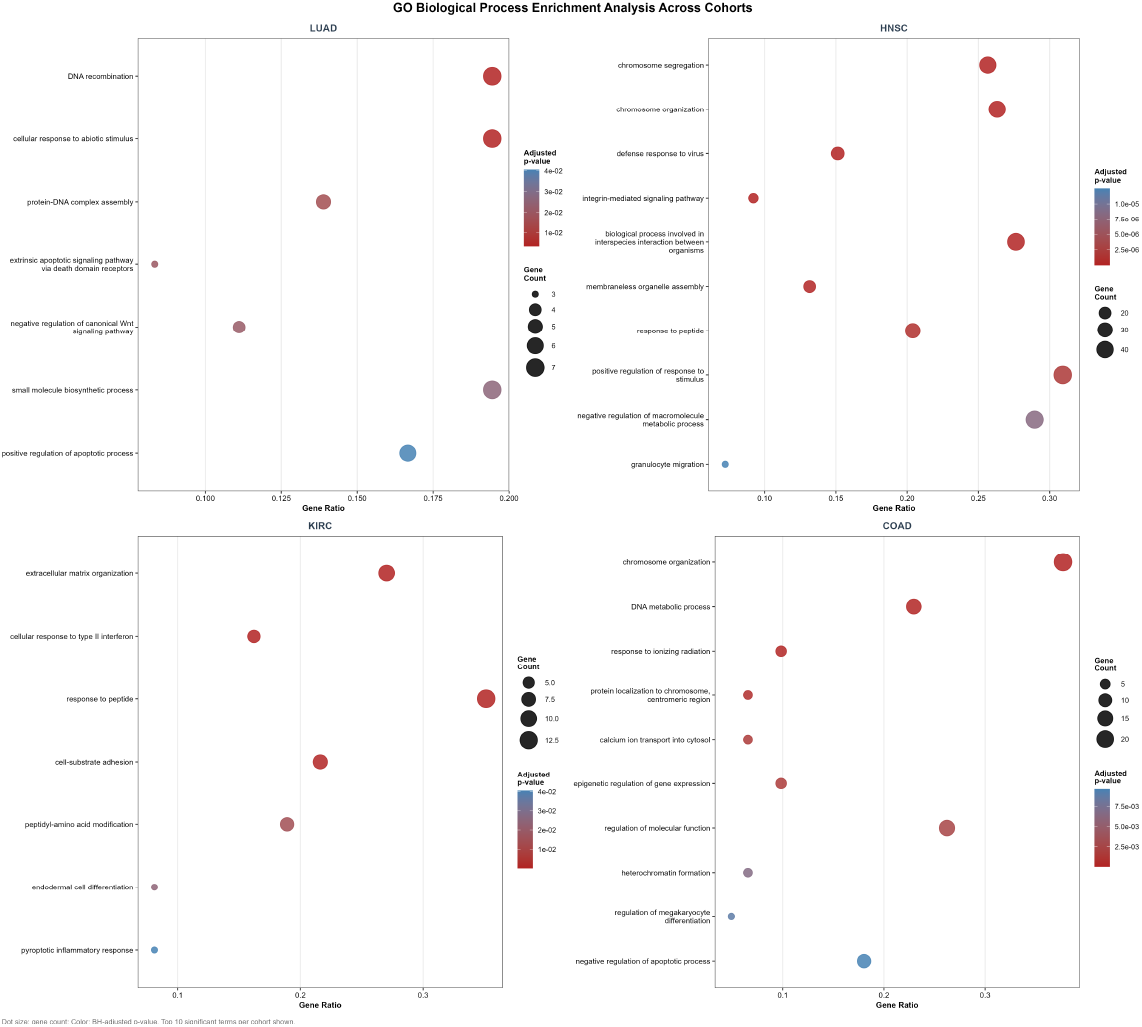
Dot plot of Gene Ontology Biological Process (GO:BP) enrichment analysis for the HR-FRGS biomarker panels across four cancer cohorts (LUAD, HNSC, KIRC, COAD). Dot size represents the gene ratio and colour intensity indicates the adjusted p-value (BH-corrected). Only terms with adjusted p < 0.05 are shown.

**Figure 9.**
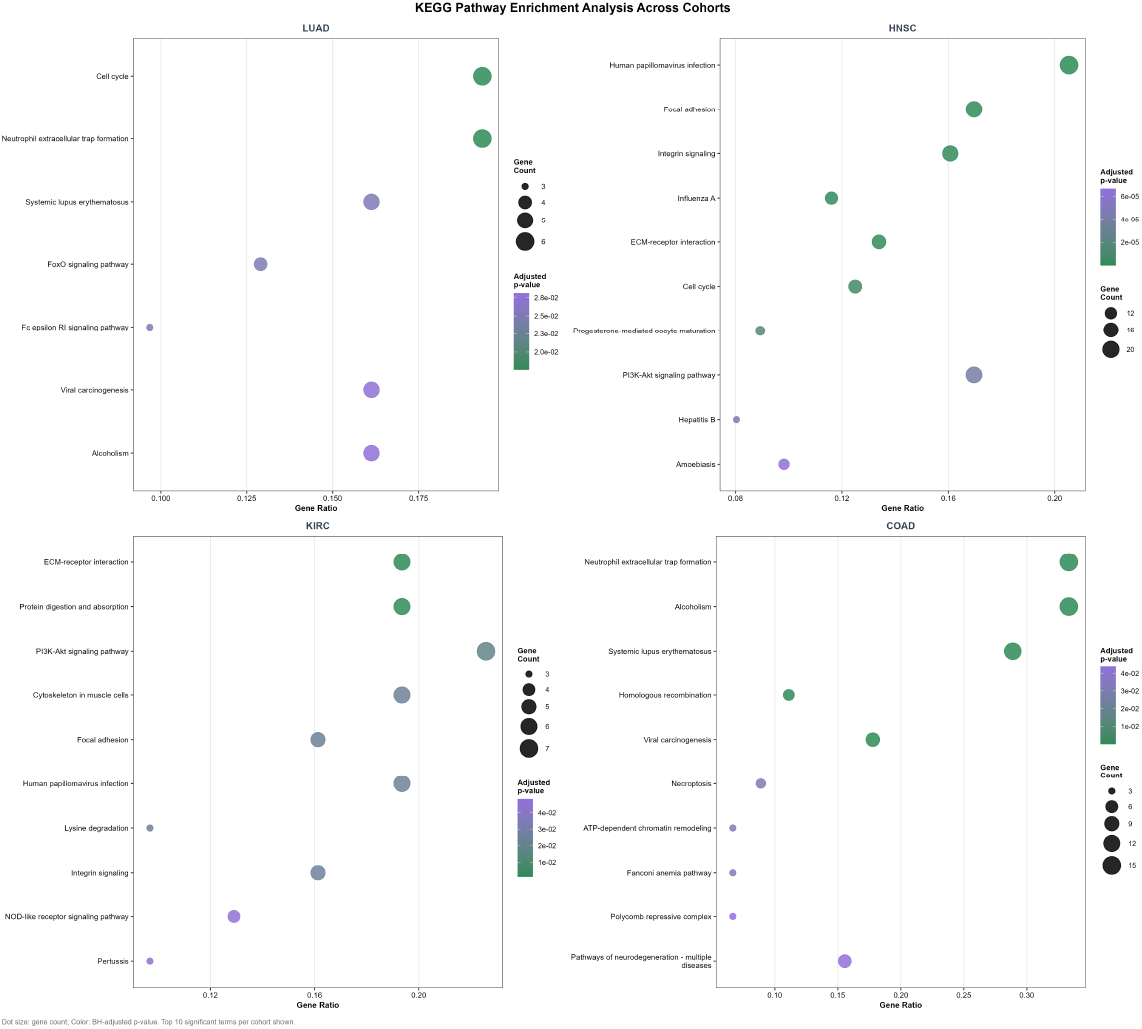
Dot plot of KEGG pathway enrichment analysis for the HR-FRGS biomarker panels across four cancer cohorts (LUAD, HNSC, KIRC, COAD). Dot size reflects the gene ratio and colour indicates statistical significance (adjusted p-value, BH-corrected). Only pathways with adjusted p < 0.05 are shown.

**Figure 10.**
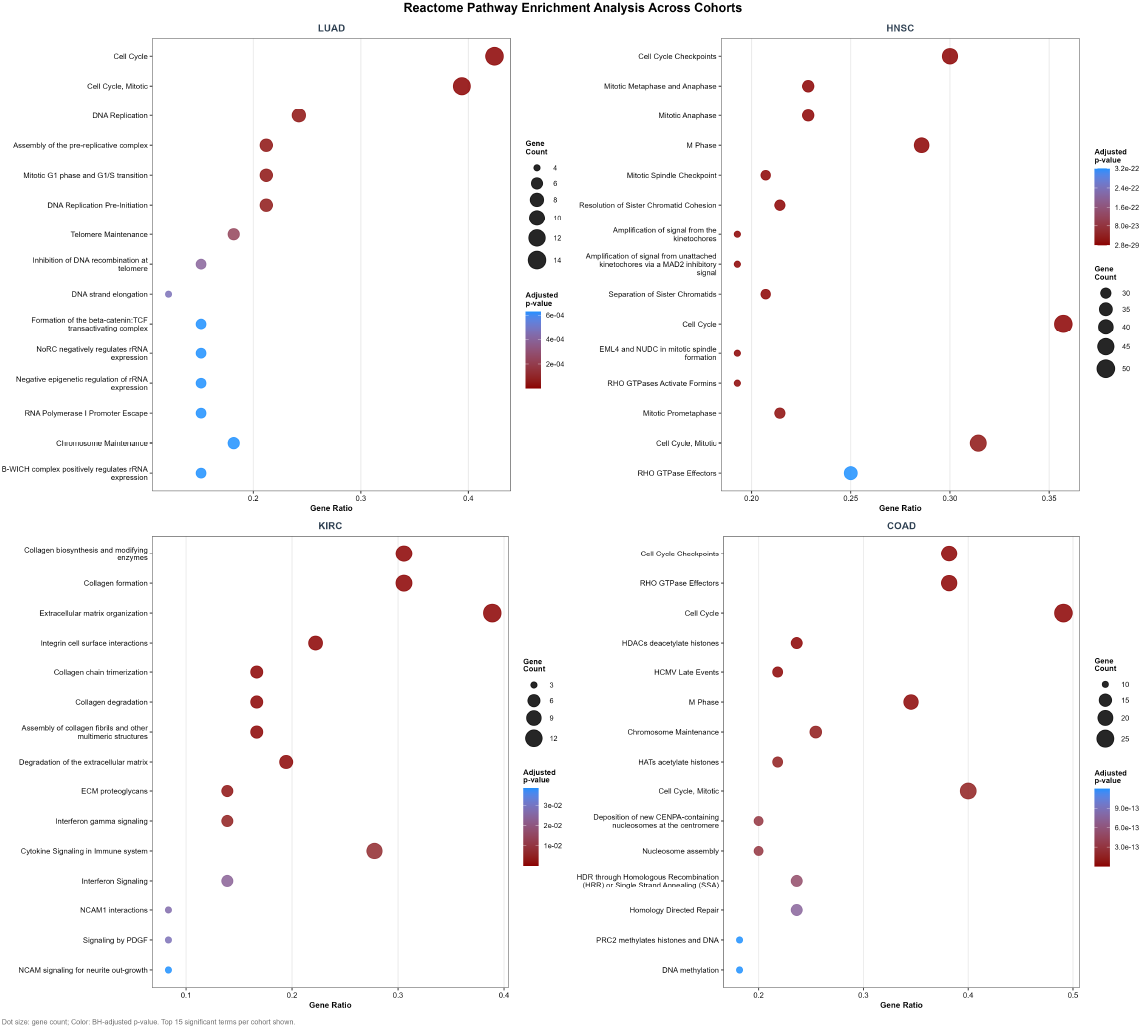
Dot plot of Reactome pathway enrichment analysis for the HR-FRGS biomarker panels across four cancer cohorts (LUAD, HNSC, KIRC, COAD). Dot size denotes gene ratio and colour encodes adjusted p-value (BH-corrected). Only terms with adjusted p < 0.05 are displayed.

Alongside shared biological context, each cohort displays unique cancer-related enrichment patterns. For example, enrichment signatures of extracellular matrix (ECM) remodelling are dominated in KIRC. In GO: BP, extracellular matrix organisation and cell-substrate adhesion and in KEGG, ECM-receptor interaction and PI3K-Akt signalling pathway enrichments show that the genes reflect the known mesenchymal character and metastasis of KIRC tumours [53]. The HNSC cohort is also enriched for ECM-receptor interaction and the PI3K-Akt signalling pathway in KEGG, alongside Human papillomavirus infection, which has the highest gene ratio among KEGG enrichments. In the GO: BP of HNSC, defence response to virus and integrin-mediated signalling pathway are significantly enriched. These results are biologically valid and expected, as HPV-driven, PI3KAkt, and integrin signalling are well-characterised drivers of HPV-associated head and neck tumours [54, 55]. LUAD is also uniquely enriched in apoptotic regulation. In GO: BP, extrinsic apoptotic signalling pathway via death domain receptors, negative regulation of canonical Wnt signalling pathway, and positive regulation of apoptotic process are specifically enriched in the LUAD cohort. The Reactome analysis also identifies the Formation of the beta-catenin: TCF transactivating complex and WNT signalling, suggesting activation of Wnt/beta-catenin signalling in lung adenocarcinoma [56]. The resulting gene sets thus contain both established and putative cancer-related genes.

## 5. Conclusion

The current study proposed HR-FRGS, a dual-layer hypergraph and fuzzy rough set framework for knowledge-driven biomarker discovery across four TCGA cancer cohorts (LUAD, HNSC, KIRC, COAD). The proposed pipeline reduces dimensionality, extracting 138 to 282 candidate biomarkers from 60,660 transcripts across all cohorts (Table 1). The candidate biomarkers also maintain or exceed the performance in internal validation compared to a much higher-dimensional gene set (DEGs, Full transcriptome). In external validation, independent GEO datasets (GSE10072, GSE75037, GSE40435) confirmed generalizability: biomarker panels achieved MCC ≥ 0.916 across all cohorts and outperformed full gene sets in XGBoost across all external datasets. The benchmarking results against the WGCNA-PPI, ML-Ensemble, and MCODE-Module pipelines showed that the candidate biomarkers from our pipeline achieve robust classification with the optimal number of genes. Also, the ablation study confirmed that removing fuzzy rough IPD-based selection and entropy filtering reduces the pipeline’s performance, especially on heterogeneous datasets such as HNSC, where Variant B MCC dropped to 0.855 from 0.908. Functional enrichment confirmed biological significance by revealing enriched pathways and processes, including cell cycle dysregulation, DNA repair, and chromatin remodelling, which have emerged as pan-cancer hallmarks (Figures 8-10). At the same time, cohort-specific signatures, such as ECM remodelling in KIRC, HPV-driven signalling in HNSC, and Wnt/*β*-catenin activation in LUAD, were also captured by the gene panel.

Despite the strong performance, several limitations in the pipeline should be acknowledged. The pipeline is totally based on STRING data with a confidence threshold of ≥ 0.7 for both hyperedge construction and OncoKB-based neighbourhood expansion. While the STRING database is curated from experimental and computational sources, the data remain biased and incomplete [33, 34]. Also, the quality of a hypergraph mainly depends on the quality of overlapping clustering. Due to noise in the STRING interaction network, incomplete PPI coverage, or sensitivity to clustering parameters, the hypergraphs may miss true higher-order biological relationships when PPI clusters are poorly formed. Although the resulting candidate biomarkers were validated computationally using an independent GEO dataset and an OncoKB gene list, no experimental validation, such as qRT-PCR confirmation of expression patterns or protein-level validation, was performed. Such wet lab validation will be necessary before these biomarkers are considered for clinical studies [2, 3]. The current study has used only bulk RNA data, which cannot account for cell-type heterogeneity in tumour microenvironments.

Our hypergraph-based approach can apply to multiomics data. miRNA expression, proteomics data, and other data types can be used as additional hyperedge layers, with each omics data type representing a different hyperedge type. This idea would enable the identification of biomarker panels that are robust across molecular modalities, reducing dependence on transcriptomics data alone [54, 57]. Applying the HR-FRGS to single-cell RNA-seq data can be valuable, as the hypergraph model can capture cell-specific regulation, and the RWR diffusion can identify genes that are central to the transcriptional programs of specific tumour subpopulations [30].

